# Autofluorescence intensity patterns encode α/β cell identity in human islets

**DOI:** 10.64898/2026.04.30.721886

**Authors:** Ivano Squiccimarro, Fabio Azzarello, Valentina De Lorenzi, Francesco Raimondi, Andrea Ghelli, Fabio Beltram, Francesco Cardarelli

## Abstract

Understanding the behavior of α- and β-cells within intact human islets is essential for elucidating mechanisms of metabolic control in diabetes. Current cell-type identification strategies rely on destructive labeling or on advanced imaging modalities such as Fluorescence Lifetime Imaging Microscopy (FLIM), which provide rich metabolic information but require specialized instrumentation and acquisition protocols. Here we show that structured intracellular intensity patterns derived from endogenous autofluorescence are sufficient to discriminate α and β cells in living human islets. Using rotation-invariant Local Ternary Pattern (LTP) descriptors combined with morphological features, we achieve highly accurate classification (AUC = 0.92), improving upon previously reported benchmarks. The resulting framework is lightweight, interpretable, and compatible with standard imaging configurations, enabling accessible and scalable analysis of label-free microscopy data. Interpretability analyses demonstrate that discrimination is driven predominantly by fine-scale intracellular intensity organization rather than global morphology. In the spectral window employed, cytoplasmic autofluorescence is prominently shaped by lipofuscin-rich granules. Consistent with prior reports of higher lipofuscin accumulation in β-cells, the dominant features identified here likely reflect differences in granule abundance and spatial organization between endocrine cell types. These findings indicate that endogenous intensity patterns encode sufficient structural information for reliable α/β discrimination, providing a biologically grounded and fully non-destructive framework for the identification of pancreatic islet cell types.

## Introduction

At the core of blood glucose regulation lies the coordinated secretion of glucagon and insulin by α and β cells within the pancreatic islets^1–4^. The activity of α and β cells is finely regulated, as alterations in their number or function can lead to severe metabolic diseases, such as diabetes^5–7^. A defining challenge in metabolic research has been observing and reliably distinguishing these cell types as they function within a living system. The current gold standard for identification, immunohistochemistry, requires cell fixation—a destructive process that affects cell viability, thereby precluding *in vivo* analyses^8–10^. Alternative strategies based on genetically encoded reporters, typically delivered through viral transduction and driven by cell-type-specific promoters, enable live-cell identification but require exogenous manipulation and may alter the native cellular state^11,12^.

The effort to visualize live islets *in vivo* has faced fundamental trade-offs. Magnetic Resonance Imaging (MRI) offers spatial resolution but suffers from low sensitivity^3,13^, while nuclear imaging (PET/SPECT) offers high sensitivity but very low spatial resolution, preventing analysis of individual islets^14,15^. Label-free optical methods offer cellular-level detail, but most (like Raman Spectroscopy or OCM) are constrained by the shallow penetration depth of light^16–18^, making them unsuitable for islet samples. Two-Photon Microscopy (2PM) overcomes this by using longer-wavelength light, enabling deep-tissue (up to 1 mm) functional imaging with low phototoxicity^19–22^. A key application, Two-Photon Fluorescence Lifetime Imaging Microscopy (2PM-FLIM), has emerged as a powerful method for non-invasive, longitudinal studies^23–25^.

In our previous FLIM-based study, we demonstrated that lifetime-derived parameters enable reliable discrimination between α and β cells (AUC = 0.86), thereby establishing a benchmark for label-free functional identification^26^. Notably, feature-importance analysis in that work revealed that a substantial portion of the classification power was already associated with static autofluorescence intensity features, with intensity-related descriptors accounting for the majority of the model contribution. In the spectral range employed (420– 460 nm under 740 nm excitation), cytoplasmic autofluorescence reflects NAD(P)H-related signals but is also prominently influenced by lipofuscin-rich granules. Previous studies, including our own analysis of the same dataset, reported a higher lipofuscin content in β-cells compared to α-cells^24,27,28^, suggesting that intensity-derived features may capture differences in lipofuscin granule abundance and spatial organization between endocrine cell types. These observations raise the possibility that spatially organized autofluorescence intensity patterns may encode α/β cell identity at the intracellular level. Despite the added metabolic information provided by fluorescence lifetime measurements, FLIM acquisition requires specialized instrumentation and extended acquisition and processing steps^29–34^, which may hinder its routine implementation in high-throughput or resource-constrained settings. Building upon these observations, the present study investigates whether the discriminative signal identified previously can be systematically recovered using intensity information alone. We show that accurate classification performance (AUC = 0.92), improving upon the previous benchmark, can be achieved using exclusively intensity-pattern and morphological descriptors, indicating that the key biological information required for α/β discrimination is robustly encoded in endogenous autofluorescence patterns. Our workflow is engineered for rotation and reflection invariance and is built on a set of intrinsic cell descriptors. These include conventional morphological features (e.g., cell area, aspect ratio, Hu moments^35^) and, critically, rotation-invariant uniform Local Ternary Pattern (LTP^36,37^) intensity-based features, which capture subcellular heterogeneity. This combined feature set enables discrimination of α and β cells using widely accessible optical configurations, avoiding specialized hardware and complex acquisition protocols (see Figure 1A). By converting standard autofluorescence imaging into a generalizable, cost-effective, and biologically interpretable tool, this work alleviates a practical limitation in live islet analysis. The presented workflow lays the foundation for high-throughput screening, real-time applications, and longitudinal studies of human islet physiology—providing an accessible avenue to investigate the cellular basis of metabolic disease.

**Figure 1.**
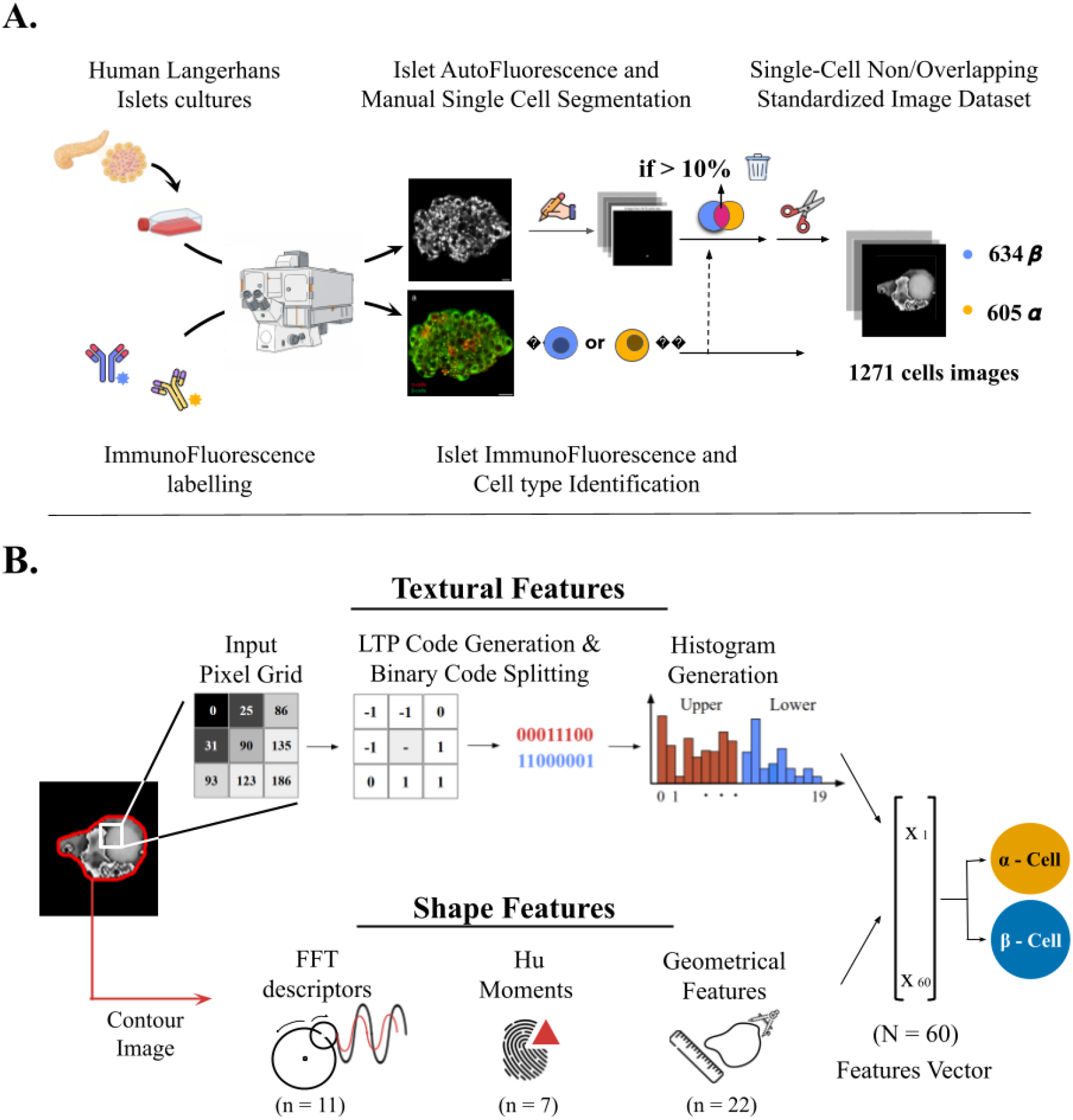
Overview of the Experimental Workflow and Proposed Pipeline. (A) Workflow from Islet Culture to Machine Learning Dataset Creation. Human Langerhans islets underwent Two-Photon Microscopy for both live-cell, label-free imaging (FLIM technique) and ground truth labeling via Immunofluorescence (IF) staining (anti-insulin (green) and anti-glucagon (red)). The autofluorescence images (top pathway) are subjected to manual single-cell segmentation. A crucial quality control step filters out overlapping cells: only single-cell images with less than 10% area overlap are retained. Retained silhouettes are cropped to a 150×150 pixel canvas, generating a standardized dataset of 1271 images (634 beta cells and 605 alpha cells) that relies solely on intensity, discarding phasor information. **(B) Feature Extraction and Machine Learning Classification:** Each standardized image is analyzed to populate a 60-dimensional Feature Vector composed of two main data types. Intensity-Pattern Features are computed using Local Ternary Pattern (LTP) codes, whose upper and lower binary components generate histograms that quantify the spatial organization of intracellular autofluorescence intensity. Shape Features quantify morphology using three approaches: FFT descriptors (boundary frequency), Hu Moments (rotationally invariant shape signatures), and general geometrical features. These descriptors are concatenated and fed into a classifier to distinguish α-cells from β-cells based solely on endogenous autofluorescence intensity.

## Results

### From image collection to dataset creation

The initial stage of the machine learning workflow builds on our previous work (Figure 1B-C) and focuses on generating a high-quality single-cell training dataset. This process comprised three main experimental phases. First, live human islets were imaged using label-free autofluorescence intensity microscopy, exciting at 740 nm and collecting emission in the 420–460 nm range, which is dominated by NAD(P)H and lipofuscin signals. Second, NAD(P)H fluorescence lifetime imaging (FLIM) was acquired at the same focal plane to capture metabolic information. Third, islets were fixed and immunostained with antibodies against glucagon and insulin to enable definitive identification of individual α and β cells.

The immunofluorescence data were spatially matched with the live-islet acquisitions, allowing precise extraction of single-cell labels from the corresponding autofluorescence images. Importantly, for the present study, the complex multidimensional FLIM datasets were deliberately reduced to total photon counts per pixel, thereby converting them into standard autofluorescence intensity images and ensuring that no lifetime information was retained. Individual cells had been manually segmented from the acquired images in the previously published dataset^38^. To prevent analytical bias arising from partially overlapping cells, any cell pair exhibiting a spatial overlap greater than 10% of the area of the smaller cell was excluded. Each retained cell was then cropped and centered within a standardized 150 × 150 pixel canvas. The final curated dataset comprised 1,271 single-cell images, including 634 β-cells and 605 α-cells.

### Exploratory Dataset Analysis

The exploration of the final dataset revealed that the classes are not trivially separable (Fig. 2). Notably, although the first two principal components capture over 56% of the dataset’s total variance (Fig.2A), the 2-D embedding of Principal Component Analysis (PCA) fails to resolve the two cell types, indicating that the primary axes of global variation in the dataset are driven by intra-class heterogeneity or other underlying factors, rather than the distinction between α and β cells (Fig.2B).

**Figure 2.**
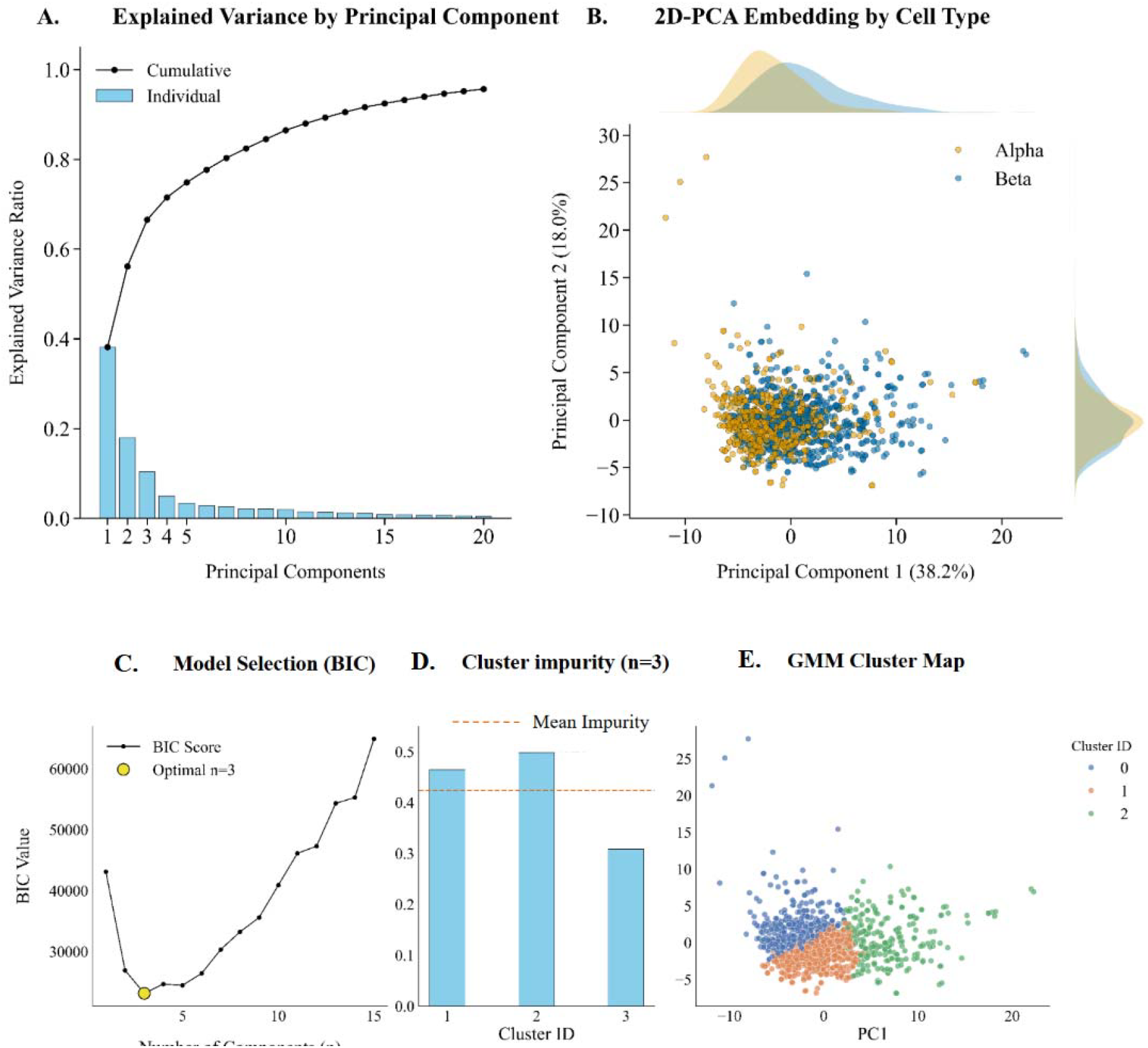
Principal Component Analysis (PCA) and Gaussian Mixture Model (GMM) Clustering of Cell Features. **(A)** Scree plot representing the individual explained variance (bars) and cumulative explained variance (line) for the first 20 Principal Components (PCs). **(B)** 2D-PCA projection onto the first two principal components (PC1 and PC2), with points colored by cell type (Alpha and Beta); marginal histograms show the distribution of cell types along each axis. **(C)** GMM model selection using the Bayesian Information Criterion (BIC) score across different numbers of components; a yellow circle highlights the optimal number of clusters (n=3). **(D)** Cluster impurity for each of the three identified GMM components, with the dashed line indicating the mean impurity (mean = 0.424). **(E)** GMM Cluster Map showing the projection of the data onto the 2D PCA space, where points are colored according to their GMM cluster assignment (0, 1, or 2).

We applied a Gaussian Mixture Model (GMM), utilizing the Bayesian Information Criterion (BIC) for model selection, to further explore the dataset’s structure. The BIC analysis identified n=3 (Fig. 2C) as the optimal number of components, demonstrating that the data’s inherent structure does not align with a simple binary classification (k=2). Rather than discrete classes, the GMM model captures a morphological continuum, identifying an intermediate “transitional” cluster that bridges the Alpha-and Beta-enriched phenotypic poles. This structural confusion, where the classes are significantly overlapped (Fig. 2E), is reflected in a high mean Gini impurity index of 0.424 (Fig. 2D) across the resulting GMM clusters and a Silhouette Score of 0.137.

A more in-depth analysis on the firsts PC1 components analysis revealed that morphological drivers, such as cell perimeter and shape complexity, evolve progressively along a continuous gradient rather than in discrete jumps, creating a biological ‘grey zone’—identified via GMM clustering as a transitional ‘bridge’—that accounts for the observed classification challenges. As detailed in the supplementary analyses, the kinetics of this phenotypic transition result in exceptionally poor linear separability (where intra-class dispersion nearly matches inter-class distance). Collectively, these data demonstrate that the alpha-to-beta transition lacks a clear low-dimensional boundary, necessitating the use of non-linear Machine Learning models to capture the underlying biological complexity (see *Supplementary Section 2*.*1*).

### Machine Learning Pipeline and Model Design

To address the non-linear separability of the data established above, we developed the machine learning pipeline outlined in Figure 3, specifically designed to ensure robustness, reproducibility, and fair model comparison. The dataset was first partitioned into training and held-out test sets using an 80/20 split, repeated across 10 independent, stratified random seeds. For each seed, nine machine learning classifiers were optimized using the training set via an inner 5-fold cross-validation scheme combined with Bayesian hyperparameter tuning (*see Supplementary Table 1*). Once optimal hyperparameters were identified, each model was retrained on the full training set, which was extensively augmented (200-fold) through random rotations and reflections to promote invariance to geometric transformations. From each cell image in both the augmented training set and the held-out test set, a 60-dimensional feature vector was extracted using a computationally lightweight process compatible with real-time applications (*see Supplementary Section S7*). During cross-validation, non-informative features exhibiting consistently low variance were pruned to improve model stability. The resulting feature vectors were assembled into a feature matrix, paired with a corresponding target vector encoding cell identity, derived from the immunofluorescence ground truth. Model performance was evaluated on the unseen held-out test set using AUC, Accuracy, Precision, Recall, and F1-score, and further assessed through stress tests involving random image rotations and reflections. To ensure reproducibility, the entire workflow was repeated independently for each of the 10 random seeds. For the final comparative analysis, models displaying excessive per-class imbalance (absolute difference > 0.2 in Precision, Recall, or F1-score) were excluded. The set of evaluated classifiers spanned linear models (Logistic Regression), kernel-based methods (Support Vector Machines), and tree-based ensembles (Random Forest, Light Gradient Boosting Machine, and XGBoost). In addition, to assess the benefit of model aggregation, we implemented meta-learning strategies, including two voting ensembles and a stacking ensemble.

**Figure 3.**
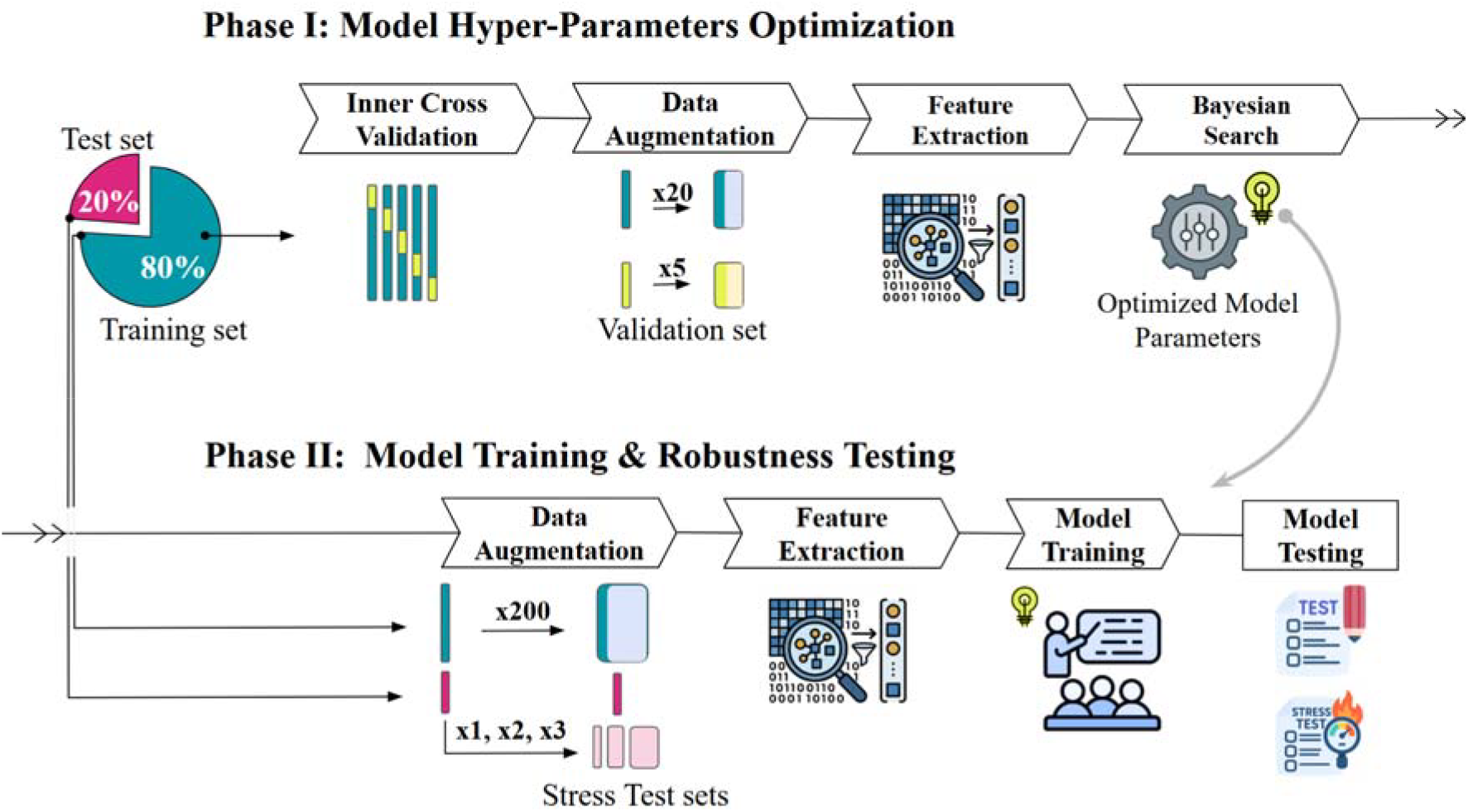
Machine Learning Pipeline for Islet Cell Type Discrimination. The workflow follows a sequential path from data partitioning to final validation across two primary phases. The process begins with the Master Dataset, which is partitioned into an 80% Training Set and a 20% Held-out Test Set across 10 random seeds (0– 9). **Phase I: Model Hyperparameter Optimization (Top)**. The training set enters an Inner 5-fold Cross-Validation loop. Within each fold, images undergo Data Augmentation (20 times for training, 5 times for validation) before Feature Extraction yields a 60-dimensional vector. These vectors drive a Bayesian Search to identify the Optimized Model Parameters (lightbulb icon) for nine distinct classifiers. **Phase II: Model Training & Robustness Testing (Bottom)**. The connection between phases is twofold: the Optimized Parameters from Phase I are fed directly into the Model Training stage, while the original 80% Training Set and 20% Test Set are passed down for final execution. Models are retrained on the augmented training data and evaluated against the unseen test set. To ensure reliability, Stress Tests are conducted using extensive augmentation on the test images. Final performance is quantified via AUC, PR-AUC, Accuracy, Precision, Recall, and F1-score.

### LightGBM Performance Surpasses Benchmarks and Is Robust Under Stress

Following performance and stability filtering, five of the nine candidate models were retained for final analysis. These models demonstrated robust performance, substantially surpassing the established benchmark (AUC 0.86) (Figure 4A). The ‘All models Voting’ ensemble achieved the highest overall metrics, with a mean Area Under the Curve (AUC) of 0.93 ± 0.02, a significant improvement over the 0.86 benchmark. This ensemble also excelled in Precision (0.87 ± 0.03) and F1-Score (0.86 ± 0.03) (Figure 4E-F).

**Figure 4.**
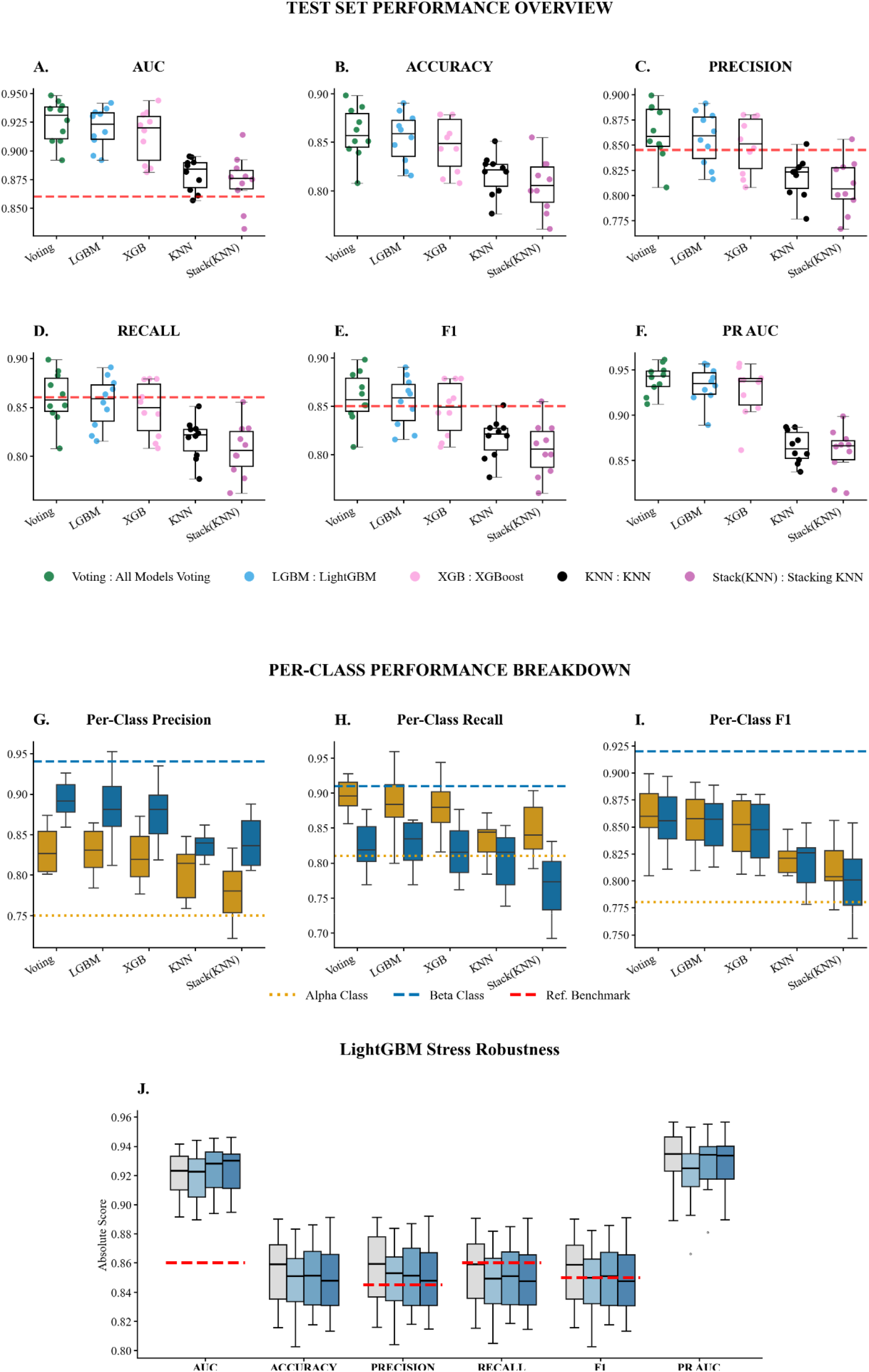
Comparative Performance of Machine Learning Models on the Test Set and Performance Stability Under Geometric Stress. This figure illustrates the classification efficacy of five machine learning models across multiple evaluation criteria. *Global Performance Metrics (A–F)*: Panels A-F show the distribution of performance scores across 10 experimental seeds for AUC **(A)**, Accuracy **(B)**, Precision **(C)**, Recall **(D)**, F1-score **(E)**, and PR AUC **(F)**. Individual data points are overlaid on boxplots to show the raw distribution, with the central horizontal line representing the median and the box defining the interquartile range (IQR). The red dashed horizontal line indicates the reference benchmark performance for each metric. *Per-Class Performance Breakdown (G–I)*: These panels provide a stratified analysis of model performance for specific cell types: Per-Class Precision **(G)**, Per-Class Recall **(H)**, and Per-Class F1 **(I)**. Performance for Alpha cells is shown in gold, while Beta cells are shown in blue. The gold dotted line and blue dashed line represent the respective benchmark performances for Alpha and Beta cell types. **(J)** Boxplots illustrate the distribution of absolute performance scores across 10 independent cross-validation seeds. Within each metric group on the x-axis, the baseline performance on pristine data (Stress Level 0) is compared against three progressively severe stress conditions. Dashed red horizontal lines indicate reference benchmark scores.

Critically, per-class metrics analysis showed that models have a balanced predictive capability between the minority (Class 0, α-cells) and majority (Class 1, β-cells) classes (Figure 4F-H). A direct statistical comparison between the ‘All models Voting’ ensemble and the best-performing *single* model, ‘LGBM’, revealed no significant difference in performance across any metric. Adhering to the principle of parsimony, the simpler LGBM model was selected for all subsequent robustness and interpretability analyses. For more information, please refer to *Supplementary Tables 2 and 3 and Supplementary Figures 5*.

The robustness of the selected Light Gradient Boosting Machine (LGBM) model was evaluated under controlled stress conditions involving random image rotations and reflections. Across all five-evaluation metrics, the model exhibited high stability, with the mean performance drop consistently remaining below one percentage point and no statistically significant differences in model performance (Figure 4J), indicating that the classifier’s predictive behavior is effectively invariant to these common geometric transformations and the robustness of the learned decision boundaries.

The largest reduction was observed for Precision (0.010 ± 0.011), whereas the Area Under the Curve remained effectively unchanged (0.000 ± 0.004). Importantly, none of the observed performance decreases were statistically significant (e.g., AUC drop, *p* = 0.938). This quantitative stability is further supported by the tight performance distributions and narrow interquartile ranges observed across all stress levels (Figure 6A). Together, these results validate the reliability of the LGBM model under realistic variations in image orientation. For more information about robustness, please refer to *Supplementary Figures 6,7 and 8 and Supplementary Tables 4-6*.

### Feature Importance and Biological Interpretation

To elucidate the predictive mechanisms underlying the Light Gradient Boosting Machine (LGBM) model, we conducted a comprehensive feature interpretability analysis combining permutation feature importance, partial dependence plots (PDPs), and Shapley Additive Explanations (SHAP). Permutation Feature Importance analysis (Figure 5A) revealed that the model’s predictive performance is predominantly driven by Local Ternary Pattern (LTP) texture features, with LTP Upper Pattern 7 (LTP Upper Pat 7 in Figure 8) emerging as the dominant predictor. These descriptors capture fine-scale subcellular texture, suggesting that intracellular spatial heterogeneity plays a central role in distinguishing cell types. While several shape-related features (specifically Hu Moments and Max Defect Depth) retained measurable importance, their contribution was markedly lower than that of the dominant texture-based feature, indicating that morphological cues alone are insufficient to fully explain the classification outcome. To further characterize the relationship between the most relevant features and the model’s predictions, Partial Dependence Plots were computed (Figure 5B). Among all examined features, LTP Upper Pattern 7 exhibited the strongest effect: the predicted probability of a cell being a β-cell increased sharply in a non-linear, sigmoidal manner as the feature value increased. This behavior identifies LTP Upper Pattern 7 as a potent intensity-pattern biomarker. Importantly, this observation aligns with the biological evidence that β-cells exhibit a higher abundance of autofluorescent granules, resulting in a more heterogeneous intracellular intensity organization that LTP-based descriptors effectively capture.. Secondary contributors, including Max Defect Depth and Inertia Eigenvalue 1, displayed qualitatively similar trends, with increasing feature values associated with a higher probability of β-cell classification, albeit with substantially smaller effect sizes. These features provide complementary information related to cell boundary complexity and mass distribution, respectively. In contrast, features such as Hu Moment 2 and LTP Upper Patterns 4 and 5 were primarily associated with the α cell class. Their PDPs exhibited flat or decreasing trends for β-cell probability, suggesting that specific global structural signatures (Hu moments) and alternative local texture patterns are more characteristic of α-cell morphology. While PDPs assume marginal independence between features, the consistency of these trends with SHAP-based analyses supports the robustness of these observations despite potential feature correlations. We performed a SHAP-based interpretability analysis aggregated across 10 independent random seeds to ensure stability and robustness of the feature attributions. The global SHAP summary plot (Figure 5C) closely mirrored the permutation importance results, confirming LTP Upper Pattern 7 as the dominant driver of model predictions. High values of this feature were uniformly associated with large positive SHAP values, strongly pushing predictions toward the β-cell class. This convergence of independent interpretability methods reinforces the conclusion that the subcellular texture captured by this LTP pattern constitutes a robust and biologically meaningful discriminator of β-cells. Conversely, secondary features such as Hu Moment 1 and Hu Moment 2 showed higher values predominantly associated with negative SHAP contributions, aligning them with α-cell identity.

**Figure 5.**
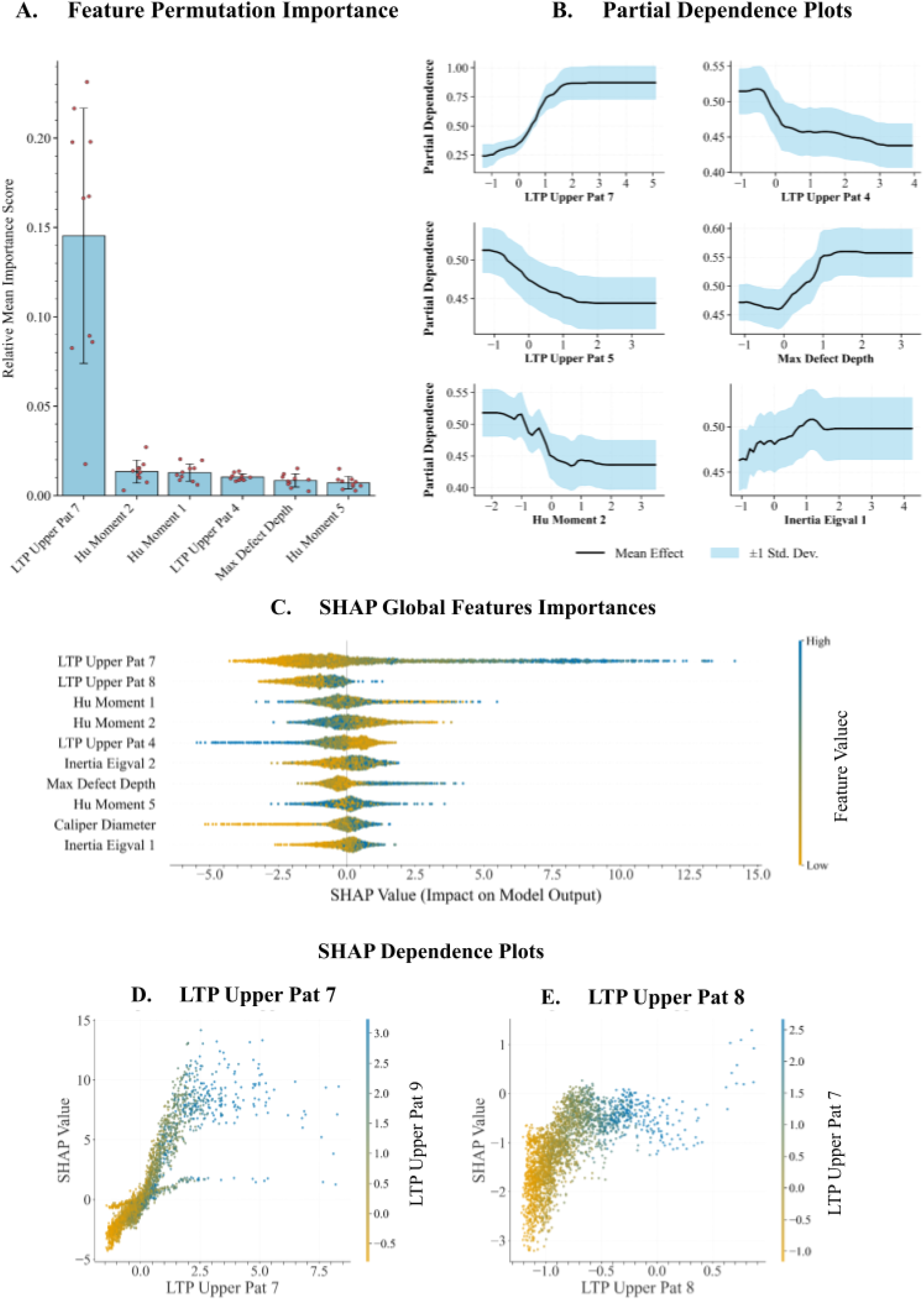
Interpretability and Feature Importance Analysis of the LightGBM Ensemble. Data aggregates results across 10 random seeds to validate biomarker robustness. **(A)** Feature Permutation Importance ranking features based on their true contribution to the model’s test set performance. Importance is quantified as the relative mean importance score (e.g., mean drop in AUC) observed when a given feature’s values are randomly shuffled across the dataset. Red dots represent the results of individual seeds. **(B)** Partial Dependence Plots for the top six critical features. These illustrate the marginal effect of a feature’s value on the predicted probability of the positive class (Class 1, hypothesized to be β cells), with the shaded area representing ±1 standard deviation across the random seeds. **(C)** SHAP summary beeswarm plot showing global feature importance ranked by mean absolute SHAP value. Dots represent individual samples, and the color denotes the feature’s value magnitude (Blue = High; Orange = Low). **(D, E)** SHAP Dependence Plots for LTP Upper Pat 7 and LTP Upper Pat 8. These plots illustrate the non-linear relationship between specific feature values and their impact on the prediction output (log-odds). The vertical color bars indicate the interaction effects with secondary features (LTP Upper Pat 9 and LTP Upper Pat 7, respectively).

Interestingly, LTP Upper Pattern 8—the second most important feature—displayed a complex, bi-modal behavior, with high feature values generally associated with positive SHAP contributions, similar to Pattern 7, but with a broader distribution of impact. Beyond individual feature effects, SHAP dependence analysis revealed that these predictors do not operate independently. As illustrated in Figure 5D, LTP Upper Pattern 7 exhibits a pronounced non-linear contribution: when its value exceeds zero, the associated SHAP value increases rapidly. The vertical dispersion of points at similar values of Pattern 7 indicates modulation by an interacting feature (LTP Upper Pattern 8), where high values of the secondary pattern (red dots) tend to cluster at the higher end of the SHAP contribution scale, amplifying the β-cell probability. A complementary relationship is observed in Figure 5E for LTP Upper Pattern 8. This feature exhibits a distinct positive trend, where increasing values correlate with higher SHAP values. The color gradient reveals a strong interaction with LTP Upper Pattern 7: observations with high Pattern 7 values (red dots) consistently yield the highest positive SHAP contributions for Pattern 8. This interaction demonstrates that the model relies on specific, reinforcing combinations of high-frequency local texture patterns, rather than isolated features alone, to achieve accurate discrimination. For more information, please refer to *Supplementary Figure 9*.

## Discussion

In this study, we developed and validated a robust and accurate machine learning framework for the label-free classification of pancreatic islet cells based solely on standard autofluorescence intensity images. We demonstrate that reliable α/β cell discrimination can be achieved using widely accessible optical setups, thereby broadening the methodological flexibility of label-free islet analysis beyond lifetime-dependent approaches. Through a rigorous comparative evaluation, a Light Gradient Boosting Machine (LGBM) model emerged as the optimal solution, achieving performance statistically equivalent to more complex ensemble approaches while adhering to the principle of parsimony. Stress testing further demonstrated that the classifier maintains stable predictive behavior under common image transformations, confirming the robustness of the learned decision boundaries and the invariance of the underlying feature representation.

Beyond classification performance, a central contribution of this work lies in its interpretability. Feature importance, partial dependence, and SHAP analyses consistently revealed that discrimination is driven predominantly by subtle intracellular intensity signatures, captured by Local Ternary Pattern (LTP) features^36,37^, rather than by global morphological descriptors. Notably, in the spectral window employed here, cytoplasmic autofluorescence includes contributions from NAD(P)H but is prominently characterized by bright lipofuscin-rich granules, which generate strong, spatially structured intensity patterns within the cytoplasm. Previous studies, including our prior work on the same dataset, showed that β-cells exhibit a significantly higher lipofuscin content compared to α-cells^27,28^. The dominant contribution of intensity-based descriptors observed here is therefore consistent with the heterogeneous spatial distribution of autofluorescent granules within the cytoplasm. In this context, the extracted intensity-pattern features likely capture differences in granule density and spatial organization, providing a quantitative and biologically grounded readout of α/β cell identity. More broadly, these findings suggest that descriptors of autofluorescence intensity patterns may act as proxies for underlying cellular states, linking image-derived features to subcellular organization and metabolic heterogeneity.

By enabling accurate cell-type discrimination from standard autofluorescence intensity images alone, the proposed framework provides an alternative implementation that does not rely on lifetime-based imaging modalities, such as FLIM. This methodological simplification substantially broadens the applicability of label-free islet analysis and opens new opportunities to:

- investigate α–β cell interactions and paracrine dysregulation without destructive labeling^8–10^;
- monitor cellular alterations during islet transplantation, including immune infiltration or early apoptotic events; and
- track functional heterogeneity within intact islets, facilitating the identification of specialized or “hub” cell populations and their network dynamics^38^.

While the present study focuses on α/β discrimination in human-derived islets, the generality of the proposed approach suggests broader applicability. Validation on independent datasets would further strengthen the generalizability of the method. However, the controlled nature of the present dataset enabled us to isolate the contribution of autofluorescence intensity patterns to cell-type discrimination with high internal consistency. Future work will be required to assess robustness across larger donor cohorts, pathological conditions, and additional endocrine cell types, as well as to integrate automated segmentation and fully end-to-end analysis pipelines.

## Materials and Methods

### Data source

All imaging data analyzed in this study were previously generated and published by Azzarello et al.^24^ and made publicly available through the associated dataset^38^. No new biological samples were collected, and no additional wet-lab experiments were performed for the present study. Details regarding human islet isolation, donor characteristics, ethical approval, two-photon microscopy acquisition, FLIM measurements, and immunofluorescence-based cell-type annotation are fully described in Azzarello et al^24^. The present work exclusively involves computational re-analysis of previously published data. Ethical approval and consent procedures were obtained in the original study^24^. Fluorescence Lifetime Imaging (FLIM) data were excluded from this analysis, which relied solely on autofluorescence intensity images.

### Dataset Curation and Preprocessing

A computational pipeline was developed to process the 1,932 raw NumPy image files and associated metadata into a curated dataset. Initially, metadata were cleaned to standardize annotations (e.g., normalizing glucose concentrations).

First, a ‘balanced’ filtering strategy was implemented: only ‘beta’ cells previously defined by F. Azzarello et al.^24^ as unsuitable for analysis were discarded. This filtering step was omitted for ‘alpha’ cells to mitigate class imbalance. A justification for the inclusion of alpha cells, along with details on the testing methodology, can be found in *Supplementary Section S1*.*3*. Corresponding results are presented in *Supplementary Section S2*.*6*.

Second, an automated overlap detection algorithm was applied to remove spatially overlapping cell masks and prevent analytical bias. Cells were grouped by experimental context and pairwise overlaps were calculated; if the shared pixel area exceeded 10% of the smaller cell’s area, both cells were discarded.

Each cell passing curation underwent a standardized processing sequence. Intensity values were normalized to an 8-bit dynamic range (0–255), and background noise was eliminated via thresholding. The primary cell contour was identified, cropped to its tightest bounding box, and resized to fit within a standardized 150×150 pixel black canvas. This fixed dimension ensured that subsequent geometric augmentations would not clip the cell image. Following the beta cell filtering process, the final dataset comprised 1,271 single-cell images of beta cells—an increase of circa 400 images compared to previous work.

### Data Augmentation

To enhance model robustness, mitigate overfitting, and address class imbalance, a multi-faceted data augmentation strategy was implemented using the OpenCV library. The augmentation pipeline relied on two fundamental geometric transformations—horizontal reflection (flip) and random rotation (0–359 degrees)—which together can generate all 2D orientations and chiral configurations of the object. A helper function randomly selected a subset of these operations to apply sequentially to each image. This process was optimized for parallel execution, ensuring diverse and robust training sets for each cross-validation fold.

### Feature Extraction

A 60-dimensional feature vector was extracted from each standardized 150×150-pixel cell image to quantify intensity-pattern and morphological characteristics. The extraction pipeline was optimized for high-throughput computation.

1. Morphological Features (40 dimensions) Shape descriptors were derived from the primary cell contour extracted using cv2.findContours. This vector comprised:
  • 22 classical morphometric features (e.g., area, perimeter, circularity);
  ▯ 7 Log-Transformed Hu Moments computed using cv2.HuMoments;
  ▯ 11 Fourier Descriptors, calculated using numpy.fft.fft, were derived by treating the contour as a complex vector. Robustness was ensured by zeroing the DC component (translation invariance), normalizing coefficients by the first fundamental frequency (scale invariance), and taking the absolute magnitude (rotation invariance).
2. Texture Features (20 dimensions) Internal autofluorescence texture was quantified using a histogram of Rotation Invariant Uniform 2 (RIU2) Local Ternary Patterns (LTP) using a custom implementation built with the standard math library. This was computed on 8-bit grayscale images with a neighborhood of P=8, radius R=1, and tolerance threshold t=5. The calculation used an optimized Numba JIT implementation with a pre-computed Look-Up Table (LUT) for instantaneous pattern-to-bin mapping.

### Model Training and Validation Strategy

The dataset was partitioned into a training set (80%) and a held-out test set (20%), stratified by cell type (using train_test_split from sklearn.model_selection). Model training and hyperparameter optimization were performed exclusively on the training partition using a 5-fold stratified cross-validation (CV) scheme (using the same package as before, StratifiedKFold).

A nested directory structure was generated to manage data splits and prevent leakage. Within each CV fold:

1. Augmentation: The training data were augmented randomly 20 times and the validation data 5 times, creating unique augmented datasets for each fold.
2. Scaling: Feature vectors were standardized using a StandardScaler. To strictly prevent leakage, this “Local Scaler” was fitted only on the augmented training features of the specific fold and subsequently applied to both the training and validation sets of that fold.
3. Feature Selection: A variance threshold (using VarianceThreshold from sklearn.feature_selection) was applied across all CV folds to identify and mask features that were consistently non-trivial (low variance), ensuring only robust features were used for modeling.
4. Outlier Detection: Possible outliers in single training sets were detected and eliminated using IsolationForest from sklearn.ensemble (contamination = 0.005).

### Hyperparameter Optimization

A Bayesian Optimization strategy using Gaussian Process regression (scikit-optimize) was employed to tune hyperparameters for each classification algorithm (e.g., RandomForest, XGBoost). The objective function maximized a combined_score, defined as the average of the Area Under the Receiver Operating Characteristic Curve (AUC) and Accuracy. The optimization ran for 40 iterations, evaluating performance via the 5-fold CV. The hyperparameter set yielding the highest mean CV score was selected. Two artifacts were saved: the optimization history and the Out-of-Fold (OOF) predictions, which were crucial for subsequent ensemble construction. For more information on the search spaces for each model, please refer to the *Supplementary Table 1*.

### Ensemble Model Construction

Three distinct ensemble strategies were developed using the OOF predictions from the cross-validation process as a meta-dataset.

▯ Greedy Selection Ensemble: An optimized voting ensemble was built using a greedy forward selection algorithm. Starting with the single best-performing model, remaining models were iteratively added if they improved the ensemble’s combined_score. The final list of selected models constituted the voting ensemble.
▯ Overall Voting Ensemble: A simple ensemble constructed by averaging the probability predictions from all candidate models.
▯ Overall Stacked Ensemble: A stacked generalization ensemble was created where OOF predictions served as the training set (X_meta) for a meta-learner. Various algorithms (Logistic Regression, Gradient Boosting Machine, K-Nearest Neighbors) were evaluated as meta-learners. The candidate yielding the highest average combined_score was selected and re-trained on the entire meta-dataset.

### Final Evaluation and Stress Testing

Optimal models, configured with the best hyperparameter combinations identified through Bayesian Optimization, were retrained on the entire training dataset after applying a 200-fold global augmentation. A Global Scaler was fitted exclusively on this augmented training set and subsequently applied to the unseen held-out test set to prevent data leakage. Outliers were detected and removed using IsolationForest (contamination = 0.005), following the same strategy applied during cross-validation.

Final model performance was evaluated on the held-out test set using AUC, PR-AUC, Accuracy, Precision, Recall, and F1-score. To assess robustness, three stress-test levels were defined by applying progressively increasing augmentation intensity to the original test set. Predictions were generated for all base models and combined using the selected ensemble strategies (Voting and Stacking), following the model-selection criteria established during cross-validation.

To ensure reproducibility, the complete pipeline was executed across 10 independent random seeds. Reported metrics represent mean ± standard deviation across seeds. Models exhibiting an absolute difference greater than 0.2 between per-class precision or recall were excluded from the final comparative analysis.

## Data Availability

The datasets generated and analyzed during the current study—including raw fluorescence intensity image frames, pre-trained ensemble models, and comprehensive interpretability results—will be made publicly available upon publication at https://doi.org/10.6084/m9.figshare.32125987. During the peer-review process, data are available from the corresponding author upon reasonable request.

## Code Availability

The complete Python pipeline, including data augmentation, feature extraction, and the model interpretability suite, is hosted on GitHub at https://github.com/ivano-squiccimarro/Autofluorescence-intensity-patterns-encode-cell-identity-in-human-islets. The repository includes instructions for reproducing the results and figures presented in this article. Access will be set to public upon formal publication of the manuscript.

## Acknowledgments

This work has received funding from the European Research Council (ERC) under the European Union’s Horizon 2020 research and innovation programme (grant agreement No 866127, project CAPTUR3D).

## Data availability statement

The authors confirm that the data supporting the findings of this study are available within the article and its supplementary materials.

## Ethics statement

The present study was exempt from requiring ethical approval

## References

1. Cabrera, O. et al. The unique cytoarchitecture of human pancreatic islets has implications for islet cell function. Proc. Natl. Acad. Sci. 103, 2334–2339 (2006).

2. Bosco, D. et al. Unique arrangement of alpha- and beta-cells in human islets of Langerhans. Diabetes 59, 1202–1210 (2010).

3. Kim, D. & Jun, H.-S. In Vivo Imaging of Transplanted Pancreatic Islets. Front. Endocrinol. 8, 382 (2018).

4. Campbell, J. E. & Newgard, C. B. Mechanisms controlling pancreatic islet cell function in insulin secretion. Nat. Rev. Mol. Cell Biol. 22, 142–158 (2021).

5. Moede, T., Leibiger, I. B. & Berggren, P.-O. Alpha cell regulation of beta cell function. Diabetologia 63, 2064–2075 (2020).

6. Kawamori, D. Exploring the molecular mechanisms underlying α- and β-cell dysfunction in diabetes. Diabetol. Int. 8, 248–256 (2017).

7. Evans, R. M. & Wei, Z. Interorgan crosstalk in pancreatic islet function and pathology. FEBS Lett. 596, 607–619 (2022).

8. Neal, A. S. et al. A method for high-throughput functional imaging of single cells within heterogeneous cell preparations. Sci. Rep. 6, 39319 (2016).

9. Baskin, D. G. A Historical Perspective on the Identification of Cell Types in Pancreatic Islets of Langerhans by Staining and Histochemical Techniques. J. Histochem. Cytochem. 63, 543–558 (2015).

10. Frikke-Schmidt, H., Arvan, P., Seeley, R. J. & Cras-Méneur, C. Improved in vivo imaging method for individual islets across the mouse pancreas reveals a heterogeneous insulin secretion response to glucose. Sci. Rep. 11, 603 (2021).

11. Jimenez-Moreno, C. et al. A Simple High Efficiency Intra-Islet Transduction Protocol Using Lentiviral Vectors. Curr. Gene Ther. 15, 436–446 (2015).

12. Shuai, H., Xu, Y., Yu, Q., Gylfe, E. & Tengholm, A. Fluorescent protein vectors for pancreatic islet cell identification in live-cell imaging. Pflüg. Arch. - Eur. J. Physiol. 468, 1765–1777 (2016).

13. Arifin, D. R. & Bulte, J. W. M. Imaging of pancreatic islet cells. Diabetes Metab. Res. Rev. 27, 761–766 (2011).

14. Wei, W., Ehlerding, E. B., Lan, X.Luo, Q.-Y. & Cai, W. Molecular imaging of β-cells: diabetes and beyond. Adv. Drug Deliv. Rev. 139, 16–31 (2019).

15. Singhal, T. et al. Pancreatic Beta Cell Mass PET Imaging and Quantification with [11C]DTBZ and [18F]FP-(+)-DTBZ in Rodent Models of Diabetes. Mol. Imaging Biol. 13, 973–984 (2011).

16. Hilderink, J. et al. Label-Free Detection of Insulin and Glucagon within Human Islets of Langerhans Using Raman Spectroscopy. PLoS ONE 8, e78148 (2013).

17. Berclaz, C. et al. Label-free fast 3D coherent imaging reveals pancreatic islet micro-vascularization and dynamic blood flow. Biomed. Opt. Express 7, 4569 (2016).

18. Lazzini, G. et al. Raman spectroscopy based diagnosis of pancreatic ductal adenocarcinoma. Sci. Rep. 15, 13240 (2025).

19. Takahashi, N. Imaging Analysis of Insulin Secretion with Two-Photon Microscopy. Biol. Pharm. Bull. 38, 656–662 (2015).

20. Li, G. et al. Multifunctional in vivo imaging of pancreatic islets during diabetes development. J. Cell Sci. 129, 2865–2875 (2016).

21. Agrawalla, B. K. et al. Two-Photon Dye Cocktail for Dual-Color 3D Imaging of Pancreatic Beta and Alpha Cells in Live Islets. J. Am. Chem. Soc. 139, 3480–3487 (2017).

22. Reissaus, C. A. et al. A Versatile, Portable Intravital Microscopy Platform for Studying Beta-cell Biology In Vivo. Sci. Rep. 9, 8449 (2019).

23. Ilegems, E. & Berggren, P.-O. The Eye as a Transplantation Site to Monitor Pancreatic Islet Cell Plasticity. Front. Endocrinol. 12, 652853 (2021).

24. Azzarello, F. et al. Single-cell imaging of α and β cell metabolic response to glucose in living human Langerhans islets. Commun. Biol. 5, 1232 (2022).

25. Gloyn, A. L. et al. Every islet matters: improving the impact of human islet research. Nat. Metab. 4, 970–977 (2022).

26. Azzarello, F. et al. Machine-learning-guided recognition of α and β cells from label-free infrared micrographs of living human islets of Langerhans. Sci. Rep. 14, 14235 (2024).

27. Cnop, M. et al. Longevity of human islet α □ and β □cells. Diabetes Obes. Metab. 13, 39–46 (2011).

28. Cnop, M. et al. The long lifespan and low turnover of human islet beta cells estimated by mathematical modelling of lipofuscin accumulation. Diabetologia 53, 321–330 (2010).

29. Mannam, V., P. Brandt, J., Smith, C. J., Yuan, X. & Howard, S. Improving fluorescence lifetime imaging microscopy phasor accuracy using convolutional neural networks. Front. Bioinforma. 3, 1335413 (2023).

30. Principles of Fluorescence Spectroscopy. (Springer US, Boston, MA, 2006). doi:10.1007/978-0-387-46312-4.

31. Datta, R., Heaster, T. M., Sharick, J. T., Gillette, A. A. & Skala, M. C. Fluorescence lifetime imaging microscopy: fundamentals and advances in instrumentation, analysis, and applications. J. Biomed. Opt. 25, 1 (2020).

32. Park, H. S. What is new in acute myeloid leukemia classification? Blood Res. 59, 15 (2024).

33. Digman, M. A., Caiolfa, V. R., Zamai, M. & Gratton, E. The Phasor Approach to Fluorescence Lifetime Imaging Analysis. Biophys. J. 94, L14–L16 (2008).

34. Becker, W. Fluorescence lifetime imaging – techniques and applications. J. Microsc. 247, 119–136 (2012).

35. Ming-Kuei Hu. Visual pattern recognition by moment invariants. IEEE Trans. Inf. Theory 8, 179–187 (1962).

36. Xiaoyang Tan & Triggs, B. Enhanced Local Texture Feature Sets for Face Recognition Under Difficult Lighting Conditions. IEEE Trans. Image Process. 19, 1635– 1650 (2010).

37. Alenizi, F. A., Mohammadi, M. & Hossein Shakoor, M. Noise-Robust Local Ternary Pattern Center for Noisy Texture Classification. IEEE Access 13, 127690–127720 (2025).

38. Azzarello, F., Cardarelli, F. & De Lorenzi, V. human Langerhans Islets data. 360626312 Bytes figshare 10.6084/M9.FIGSHARE.23765169.V1 (2023).

39. Benninger, R. K. P. & Hodson, D. J. New Understanding of β-Cell Heterogeneity and In Situ Islet Function. Diabetes 67, 537–547 (2018).

