## Supplementary information and figures for "Autofluorescence intensity patterns encode α/β cell identity in human islets"

**S1 Implementation Details**

**S1.1 Models Search Spaces Summary**

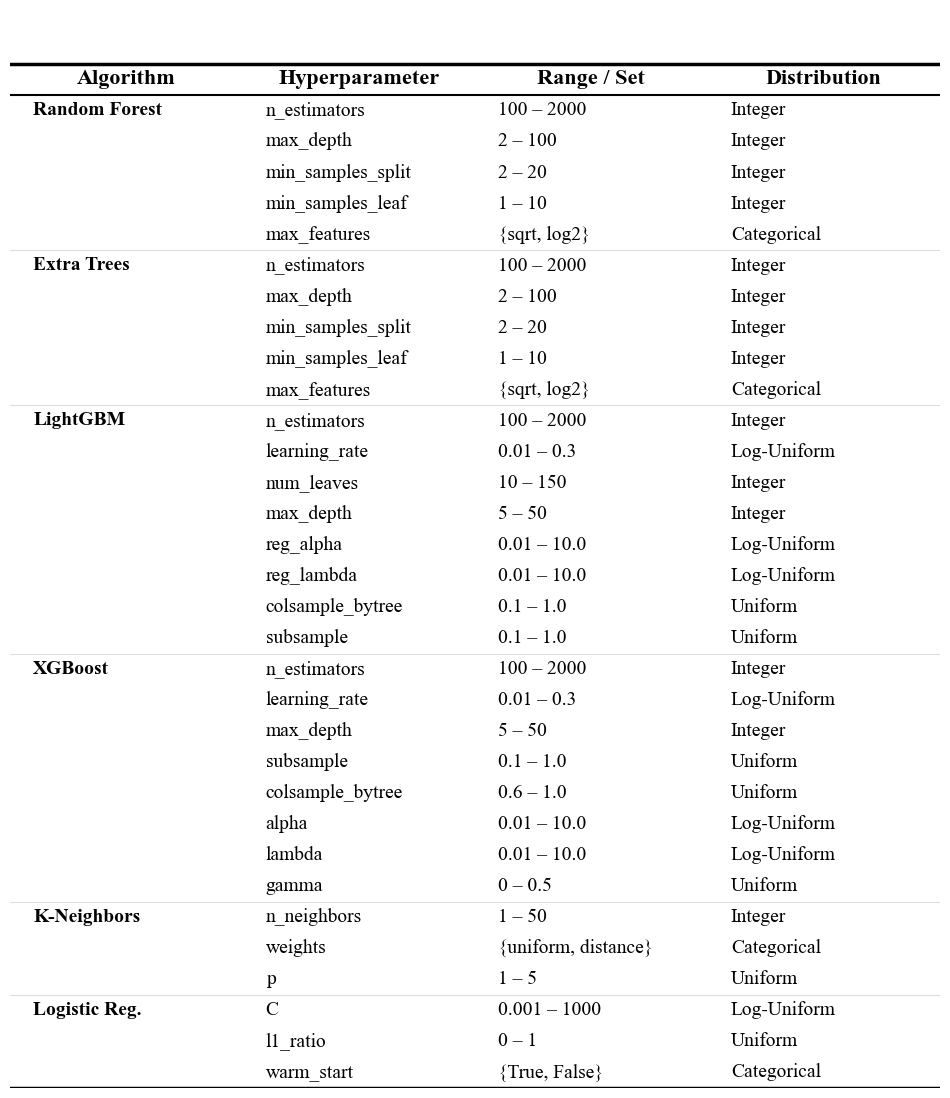
**Supplementary Table S1| Hyperparameter Search Spaces.** *The table details the range and distribution of hyperparameters optimized via Bayesian Optimization for each base classifier. "Log-Uniform" indicates the search was performed on a logarithmic scale.*

**Note on Stacking Meta-Learners:** The Stacking Ensemble evaluated three candidate meta-learners: Logistic Regression (max iterations: 5000), K-Nearest Neighbors, and Gradient Boosting Classifier. These were evaluated using their default hyperparameters (scikit-learn defaults) to select the best architecture for the meta-layer.

**S1.2 Speed Benchmarking Pipeline**

We evaluated model performance through a structured three-phase benchmarking approach comprising computational profiling, scalability analysis, and real-time system validation.

First, we assessed the baseline computational efficiency by measuring per-image latency and throughput for both the original serial pipeline (n_jobs=1) and the parallelized engine (n_jobs=-1) using the LightGBM classifier. To ensure robust statistical significance, we employed a strict protocol: 5 warm-up runs to mitigate initialization overhead, followed by 50 measurement repeats using high-resolution performance counters. The evaluation dataset was expanded via random augmentation (N=10 per image) to minimize timer variance. Results were visualized via violin plots to capture the underlying density distribution of processing times for Feature Extraction and Model Prediction stages.

Subsequently, system scalability was evaluated across a logarithmic range of batch sizes (N=10^1 to 10^5). The dataset was dynamically augmented to meet target volumes, allowing us to characterize the transition from initialization-bound execution to steady-state throughput saturation for both execution modes.

To evaluate real-time feasibility under single-frame conditions, we optimized the inference pipeline to minimize per-frame overhead and isolate algorithmic latency from hardware parallelism. A representative subset of high-performing LightGBM models was selected based on test-set AUC and prediction stability across random seeds. The optimized inference engine was benchmarked against the standard research pipeline using a standardized stream of N = 2,000 augmented frames over five independent repetitions. Median latency, throughput (FPS), and prediction fidelity were recorded. Predictive consistency between implementations was verified using Pearson correlation, mean absolute error (MAE), and label agreement.

**S1.3 Validation of Extended Alpha Cell Inclusion and Model Robustness**

To validate the discriminative contribution of the reintroduced alpha cell images (Dataset++) and distinguish their informational content from random noise, we conducted a comparative analysis using LightGBM classifiers trained on three distinct data configurations: the extended dataset (Dataset++), the original restricted dataset (Dataset), and the restricted dataset balanced via Synthetic Minority Over-sampling Technique (Dataset+SMOTE). To ensure statistical reliability, all experiments were replicated across 10 random seeds.

The evaluation methodology consisted of two analytical strategies. First, we performed an Inter-Dataset Generalization Analysis to assess performance on unseen, independent data partitions. This involved cross-evaluating models trained on the restricted splits against unique samples from the extended splits, and vice versa, to quantify generalizability beyond the training distribution. Second, we conducted an Intra-Dataset Robustness Analysis (Stress Test) to isolate the genuine signal from the synthetic artifacts. Models were evaluated on the standard test set and subsequently subjected to high-intensity noise injection (Level 3) to monitor the stability of class-specific decision boundaries (specifically Alpha vs. Beta precision shifts). Performance was quantified using Accuracy, AUC, F1 Score, and per-class Precision/Recall. Statistical significance for pairwise comparisons was assessed using the Wilcoxon Signed-Rank Test with a significance threshold of 𝛼= 0.01.

**S2. Results**

**S2.1 Exploratory Dataset Analysis and Machine Learning Approach Justification**

At first, in continuation with previous work, we applied the K-means clustering algorithm to assess the intrinsic separability of the dataset independently of any model assumptions regarding class labels (Supplementary Fig. S1).

To evaluate the optimal number of clusters (*k*), we examined the Within-Cluster Sum of Squares (WCSS) using the elbow method. As shown in the WCSS curve (Supplementary Fig. S1A), no clear elbow point corresponding to the true number of biological classes (*k* = 2) was observed. Instead, the WCSS decreased gradually with increasing *k*, indicating the absence of a natural low-dimensional binary clustering structure. To investigate whether higher values of *k* could capture local variance and meaningful class separation, a representative value of *k* = 10 was selected (Supplementary Fig. S1A).

To quantitatively assess cluster purity, we computed the Gini impurity index for each of these 10 clusters using the ground-truth labels. In this context, a Gini index of 0 corresponds to a perfectly homogeneous cluster, whereas a value of 0.5 indicates maximal (50/50) class mixing. The resulting analysis (Supplementary Fig. S1B) demonstrated that none of the clusters approached purity. The elevated mean Gini impurity of 0.375 across the subgroups further confirmed that α-cell and β-cell samples remain highly intermingled even when artificially partitioned into granular clusters (Supplementary Fig. S1C).

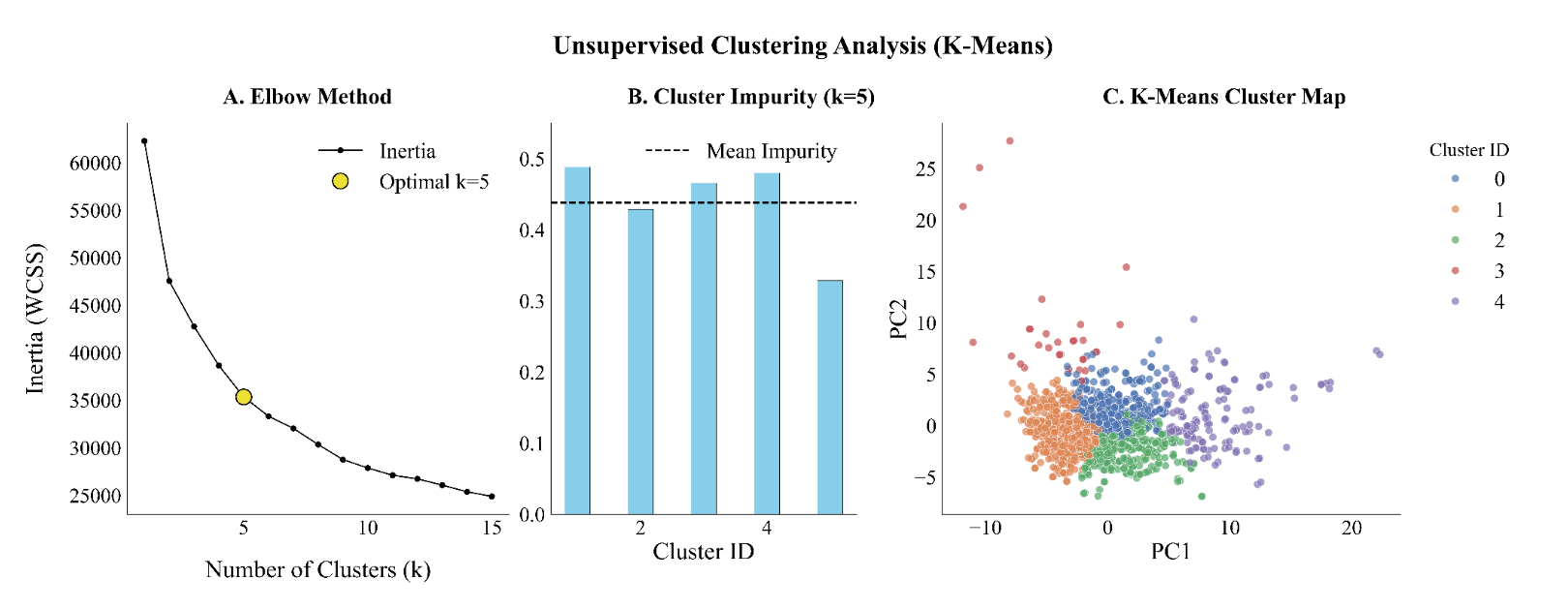
 **Supplementary Figure 1 | Unsupervised Clustering Analysis of Cell Features. (A)** **Elbow method analysis using Within-Cluster Sum of Squares (WCSS)** to determine the optimal number of clusters (k). The orange star indicates the selection of k=10 as the optimal cluster count, marking the point where the rate of WCSS reduction diminishes significantly. **(B)** The **Gini Impurity Index** was calculated for each of the 10 identified clusters to assess cluster homogeneity. The dashed orange line represents the mean Gini Index (0.375) across all clusters. **(C)** Visualization of the 10 identified clusters projected onto the first two Principal Components (PC1 and PC2). Individual cells are colored according to their assigned Cluster ID (0–9), illustrating the spatial distribution and separation of cell subpopulations in the reduced feature space.

Building upon the limitations of rigid, distance-based algorithms, we subsequently applied a Gaussian Mixture Model (GMM) to better accommodate the complex, overlapping distributions of the data. As noted in the main article, the Bayesian Information Criterion (BIC) identified *n = 3* components as the optimal fit.

Rather than representing a classification error, the GMM clustering reveals that this three-cluster structure successfully captures a continuous morphological gradient. The GMM identifies a distinct intermediate "transitional" cluster that acts as a biological bridge between the α-enriched and β-enriched phenotypic poles (Supplementary Figure S2).

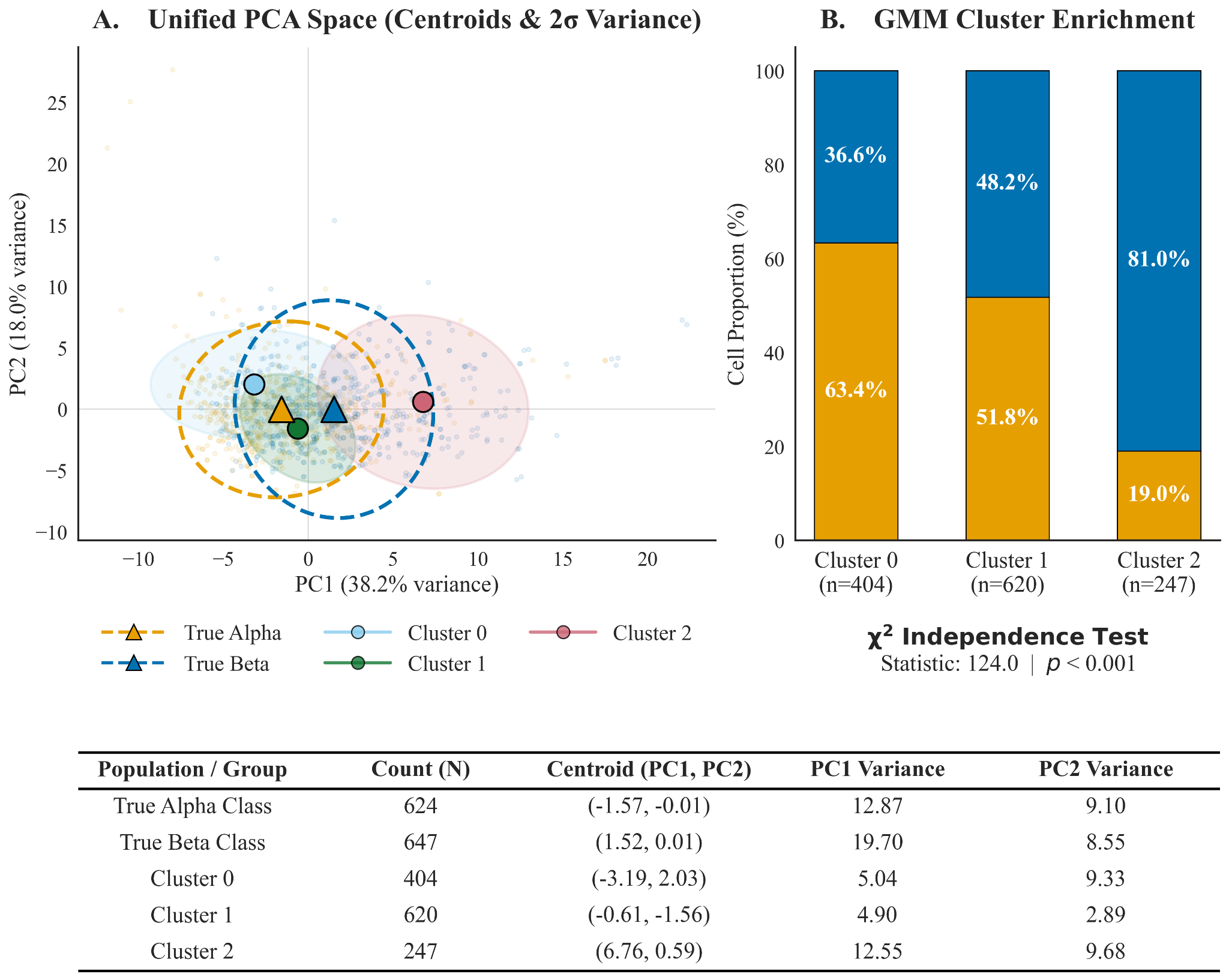
**Supplementary Figure S2 | Unified PCA Space and Cluster Enrichment**. (Left) Principal Component Analysis (PCA) projecting both the true biological classes (dashed ellipses, triangle centroids) and the optimal *n=3* GMM clusters (solid ellipses, circular centroids). The true Alpha and Beta classes exhibit massive structural overlap. In contrast, the GMM clusters map a clear trajectory across the principal components, with Cluster 1 occupying the highly mixed center. (Right) Stacked bar chart and $\mathbf{\chi}^{2}$ independence test (*p < 0.001*) confirming cluster enrichment. While Clusters 0 and 2 are heavily enriched for Alpha (63.4%) and Beta (81.0%) cells, respectively, Cluster 1 is highly heterogeneous (51.8% Alpha, 48.2% Beta). (Bottom) The accompanying statistical table highlights the high PC1 and PC2 variances for the true classes, contrasting with the tighter, more localized variances of the unsupervised GMM clusters, further emphasizing the dataset's dense central overlap.

By mapping the primary feature drivers along this PCA trajectory, we observed that parameters such as cell size (Perimeter) and macroscopic shape (Caliper Diameter, Major Axis) evolve progressively rather than in discrete jumps (Supplementary Figure S3).

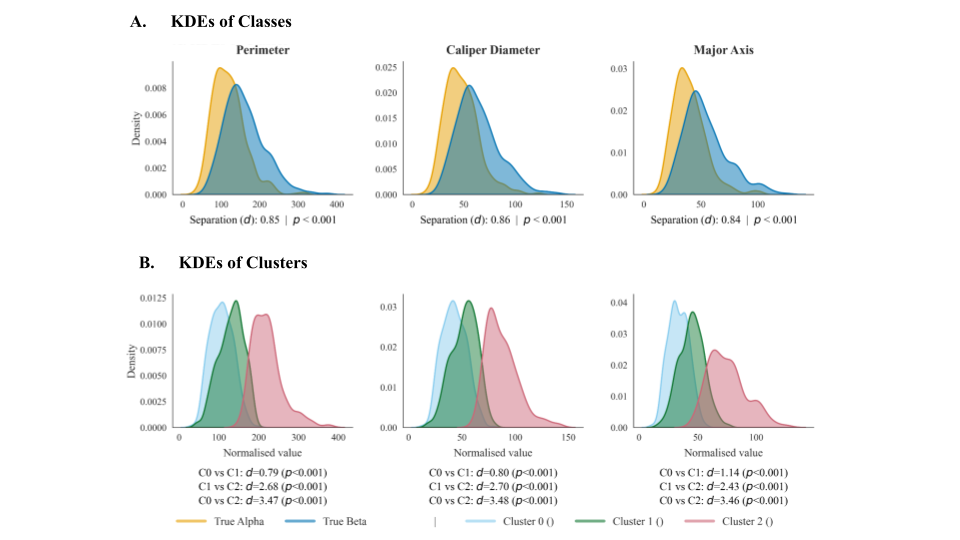

**Supplementary Figure S3 | Morphological Gradient KDEs and Separation Metrics. Kernel Density Estimates (KDEs) for the top PC1-driver features**. (Row A) Distributions of the true Alpha and Beta classes. While statistically significant (*p < 0.001*), the structural overlap is substantial, yielding a moderate Cohen’s d separation of ~0.85 across size features. (Row B) The same features partitioned by the *n=3* GMM clusters, quantitatively proving the existence of a continuous gradient. The separation between the extreme poles (Cluster 0 vs. Cluster 2) is massive (*d > 3.45*). However, the separation between adjacent clusters (e.g., C0 vs C1, circa 0.80) mimics the true class separation, proving that Cluster 1 acts as a biological "stepping stone" or transitional state rather than a distinct outlier.

This non-trivial separability and the continuous kinetics of the phenotypic transition perfectly characterize the biological "grey zone" in the dataset, explaining why simple clustering approaches struggle to isolate the two cell types.

Across numerous morphological features, we observe this continuous α-to-β transition rather than a clean, discrete boundary. Attempting to force a single, optimal dividing line through this continuum to separate the data essentially mimics a Linear Discriminant Analysis (LDA) approach (Supplementary Figure 4). However, our dataset demonstrates exceptionally poor linear separability: the overlap between α-cell and β-cell feature distributions is reflected in a low Fisher’s Ratio of 1.226, indicating limited between-class variance relative to within-class variance. This observation is further corroborated by a geometric Separation Ratio of just 1.06, where the mean inter-class distance (9.20) only marginally exceeds the average intra-class dispersion (8.70).

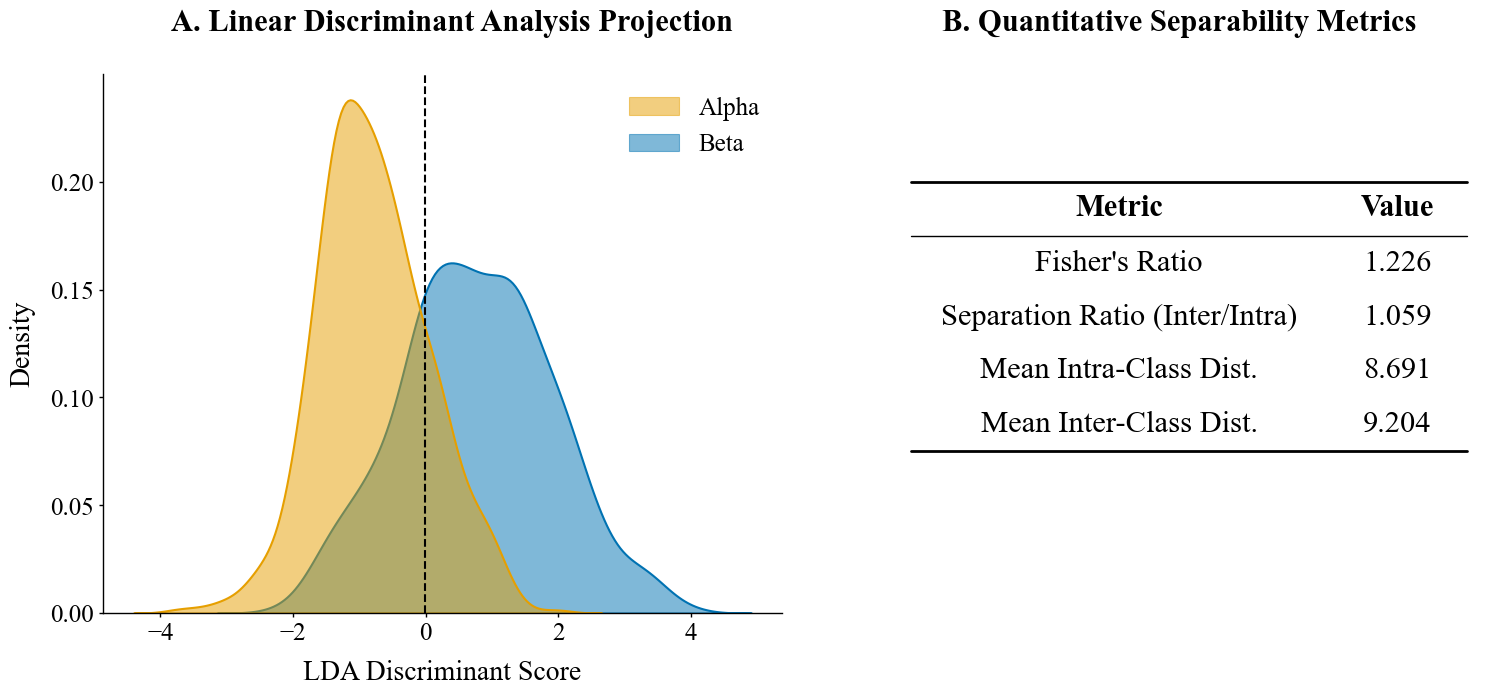
**Supplementary Figure 4 | Linear Separability Assessment via LDA Projection.** This figure evaluates the linear separability of Alpha and Beta cell features by projecting them onto the optimal discriminant axis identified by Linear Discriminant Analysis (LDA). **(A)** LDA Projection Density. Kernel Density Estimation (KDE) plots visualize the distribution of Alpha (red) and Beta (green) cell features along the single LDA component that maximizes class separation. The substantial overlap between the two distributions visually demonstrates the difficulty of separating these classes using a linear boundary. The vertical dashed line represents the optimal linear decision boundary. **(B)** Quantitative Separability Metrics. A summary table provides statistical metrics quantifying the separation. The low Fisher's Discriminant Ratio confirms the significant overlap observed in the density plots. Additional metrics, including the separation ratio and mean inter/intra-class distances, further substantiate that the feature space is not linearly separable, justifying the necessity for non-linear classification models.

Collectively, the unsupervised clustering results, the prominent transitional "grey zone," and the failure of classical linear separability metrics demonstrate that the dataset is not linearly separable and lacks a clear low-dimensional partitioning structure. These characteristics provide compelling empirical justification for the adoption of non-linear Machine Learning models, which are uniquely suited to capture the complex and subtle morphological patterns required for accurate α-cell and β-cell discrimination.

**S2.2 Models Performances and Principle of Parsimony**

As stated in the main article, some models demonstrated the ability to substantially surpass the established benchmark and to achieve a good balance between precision and recall for both classes (see Supplementary Table 2).

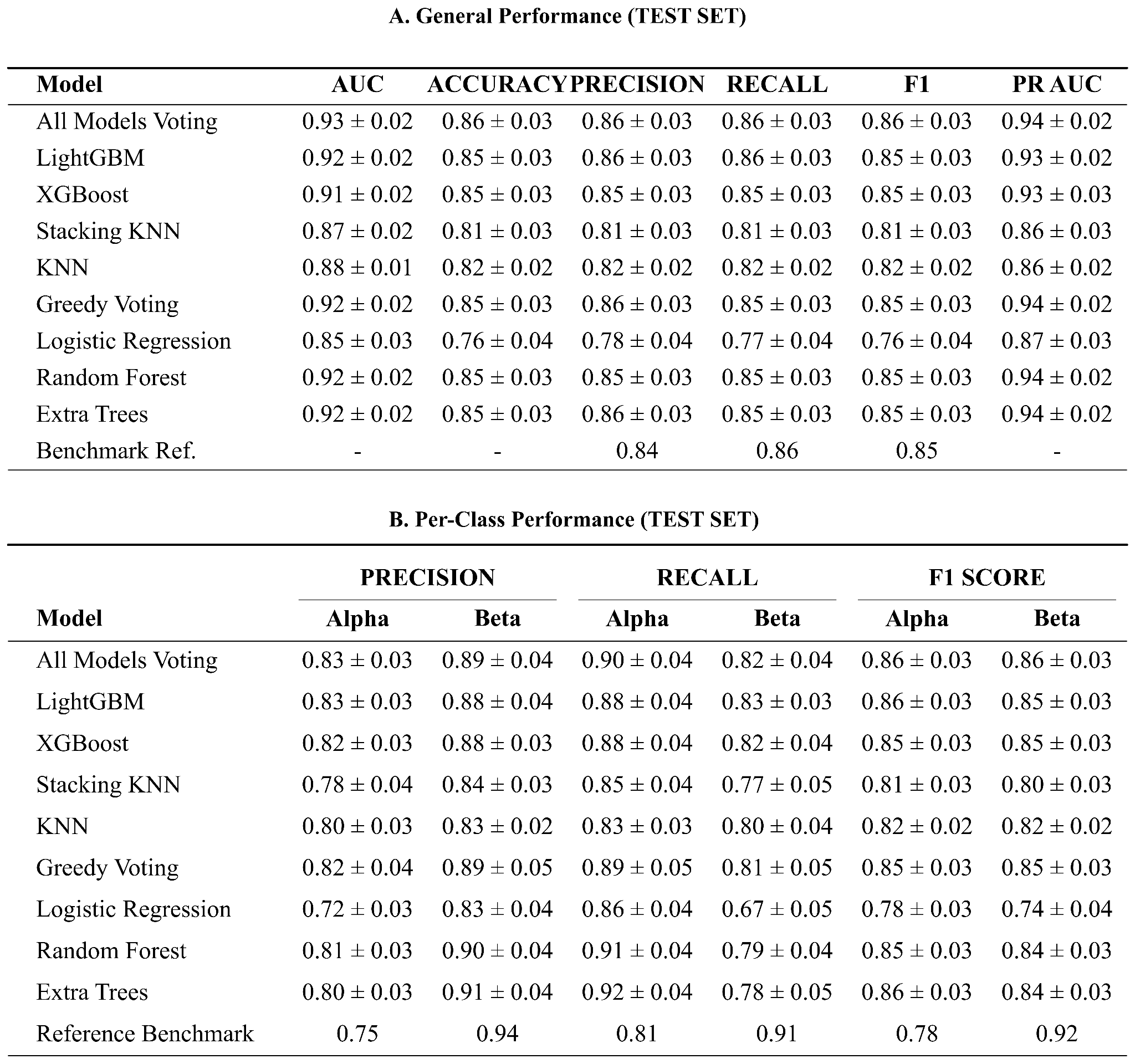

**Supplementary Tables 2 | Quantitative Performance Summary of Machine Learning Models on the Test Set.** These tables provide a detailed numerical evaluation of the five distinct machine learning models compared against a reference benchmark, shown in Figure 5 in section 2.3, LGBM Model Performance Surpasses Benchmarks of the main article. **(A) Global Test Set Results.** This table summarizes the overall performance metrics (AUC, Accuracy, F1-Score, Precision, and Recall). Data is presented as the mean ± standard deviation across experimental runs. The "All models Voting" and "LGBM" classifiers demonstrate top-tier performance, consistently outperforming the Reference values across all key metrics. **(B) Per-Class Metrics.** This table breaks down the performance by specific cell type, detailing Precision, Recall, and F1-Scores for both Alpha and Beta cell populations. This granular view highlights the models' ability to correctly identify and distinguish between the two specific classes relative to the reference standards.

Despite combining the models into an All Voting Ensemble, the resulting performance improvements were not statistically significant when compared to the top-performing individual model, LGBM (Mann-Whitney U-test, p > 0.05). For example, the Area Under the Curve (AUC) for the ensemble (0.93 +/- 0.2) remained statistically indistinguishable from the LGBM classifier (0.92 +/- 0.02), p = 0.521 (see Supplementary Figure 5 and Supplementary Table 3). The same can be said for all other metrics.

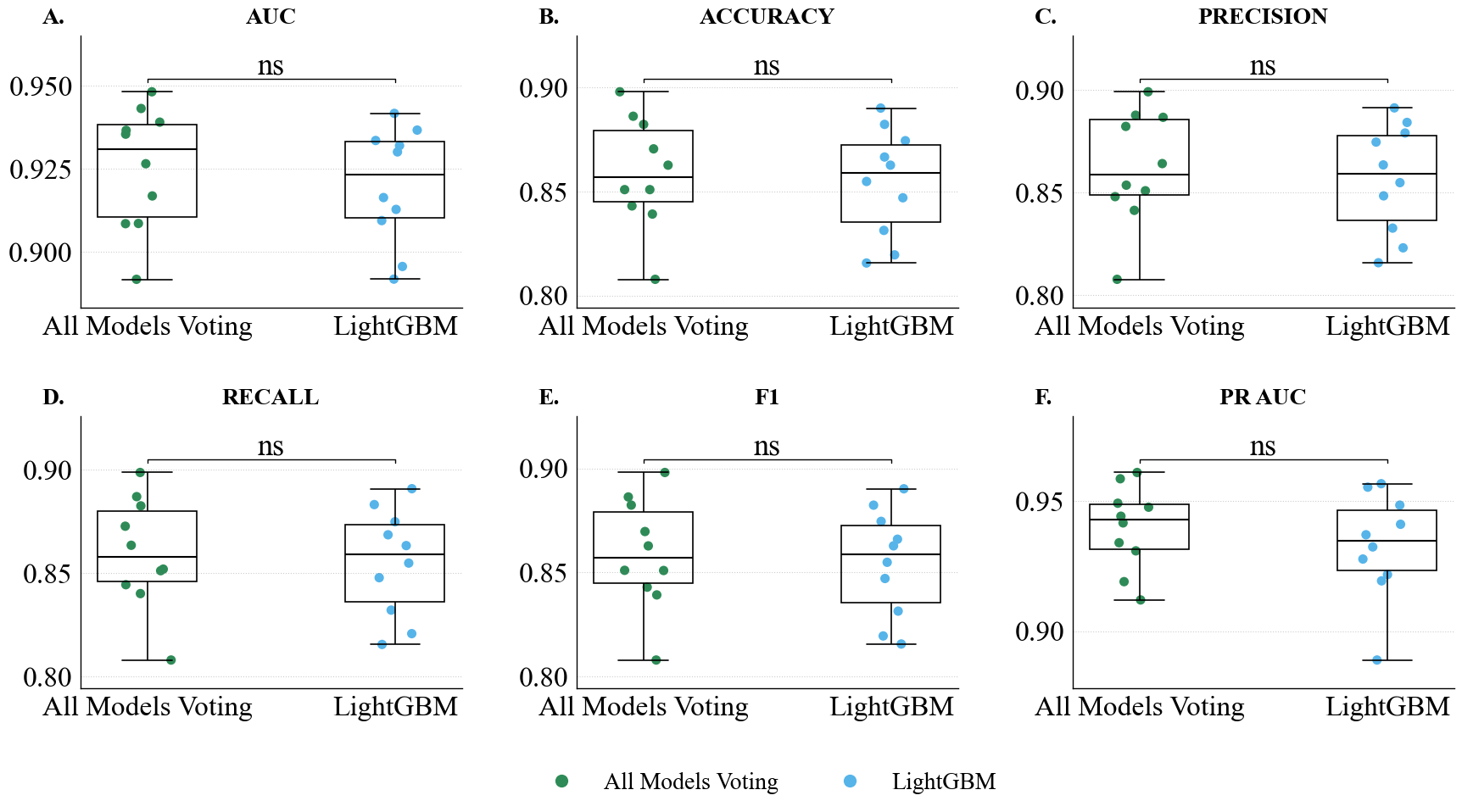

**Supplementary Figure 3 | Statistical Comparison of the Top-Performing 'All models Voting' Ensemble and the 'LGBM' Model.** The figure provides a direct performance comparison between the 'All models Voting' ensemble and the individual 'LGBM' model to determine if a statistically significant difference exists between them. **A–E** boxplots in the top panel visually compare the performance distributions of the two models across five key metrics. While the 'All models Voting' ensemble consistently exhibits slightly higher median performance, there is substantial overlap in the interquartile ranges, suggesting that the performance difference may not be statistically meaningful.

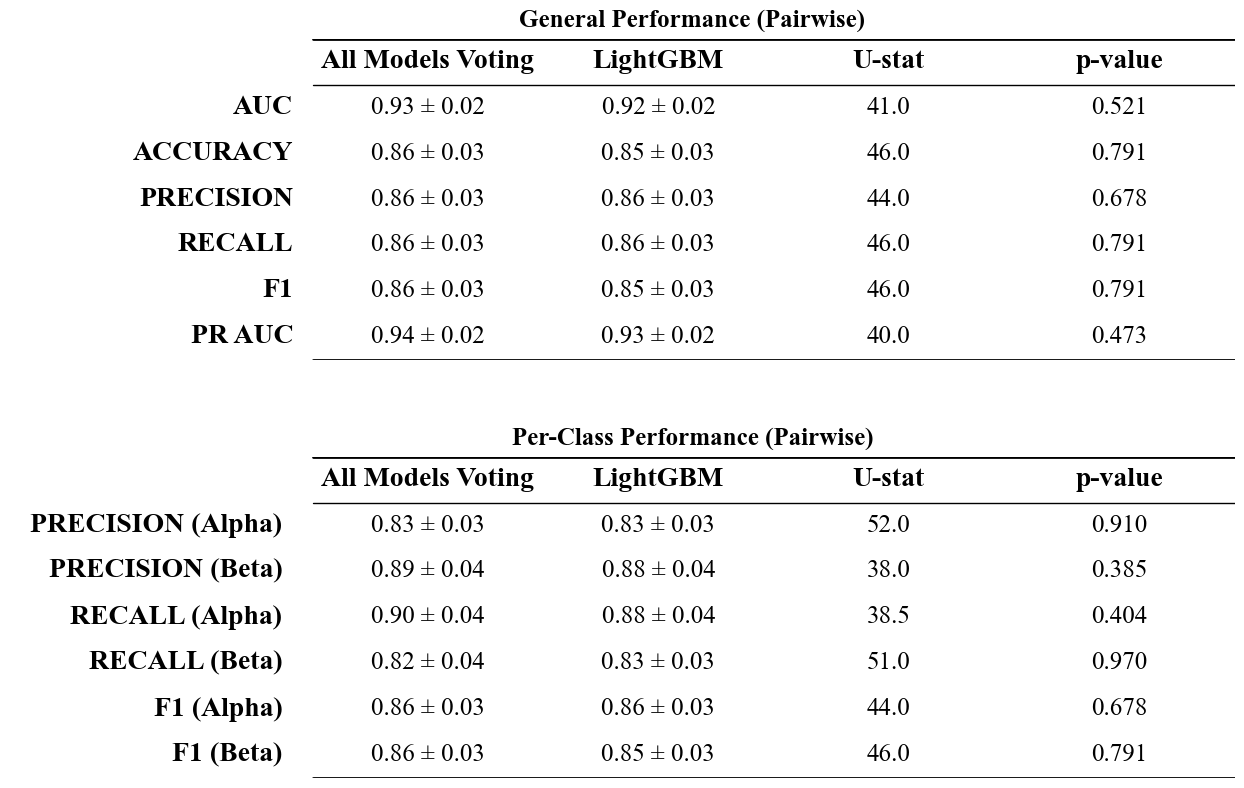
**Supplementary Table 3 | Statistical Comparison of Top-Performing Models.** This table presents a pairwise statistical comparison between the "All models Voting" ensemble and the individual "LGBM" model using the Mann-Whitney U test. The mean performance scores (± standard deviation) are listed for five key metrics: AUC, Accuracy, Precision, Recall, and F1-Score. The "U-statistic" and "p-value" columns quantify the statistical significance of the performance differences. A p-value greater than 0.05 across all metrics indicates that there is no statistically significant difference in performance between the ensemble method and the standalone LGBM model.

**S2.3 Models Robustness on Stress**

As stated in *Section 2.4 LGBM Model Performance Surpasses Benchmarks and Is Robust Under Stress,* the main article, the robustness of the selected LGBM model to rotational and reflective perturbations was quantitatively assessed. Across all evaluated metrics, performance variations remained minimal and statistically non-significant, indicating stable behavior under the applied stress conditions (see Supplementary Table 4).

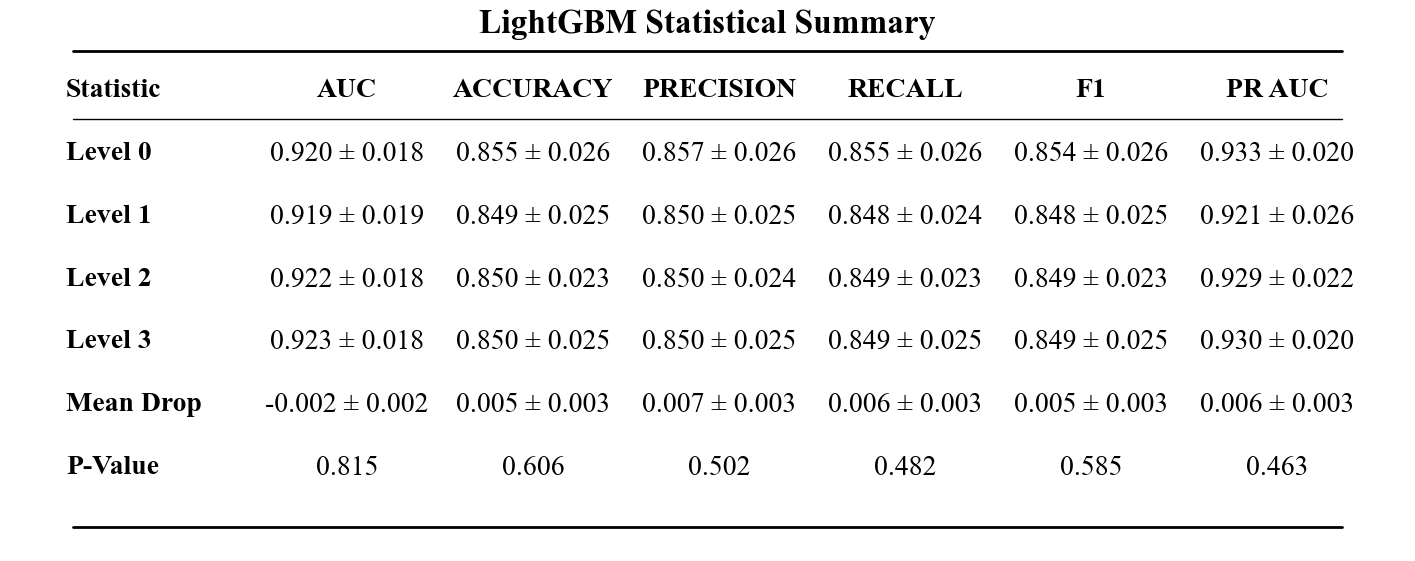

**Supplementary Table 4 | Detailed Statistical Summary of LGBM Model Performance and Equivalence Under Geometric Stress.** Absolute performance metrics (AUC, Accuracy, Precision, Recall, F1-score, and PR AUC) for both the pristine baseline (Level 0) and subsequent stress conditions (Levels 1–3) are reported as the Mean ± Standard Deviation (SD) to illustrate cross-validation variance across the 10 random seeds. To rigorously evaluate model robustness, the overall performance degradation is quantified as the Mean Drop (Baseline minus Stress) and is reported alongside its 95% Confidence Interval (CI) to indicate statistical precision. The mean drops approach zero and are tightly bound by their CIs. Furthermore, nonparametric Mann–Whitney U testing confirms no statistically significant difference between the pristine and stressed distributions across any metric (all p > 0.05).

What we previously said for LGBM is also valid for the other models ( as detailed in Supplementary Figures 6-7 and Supplementary Table 5-6): the robustness of the constructed features made the models' predictions and performance invariant to rotation and reflection.

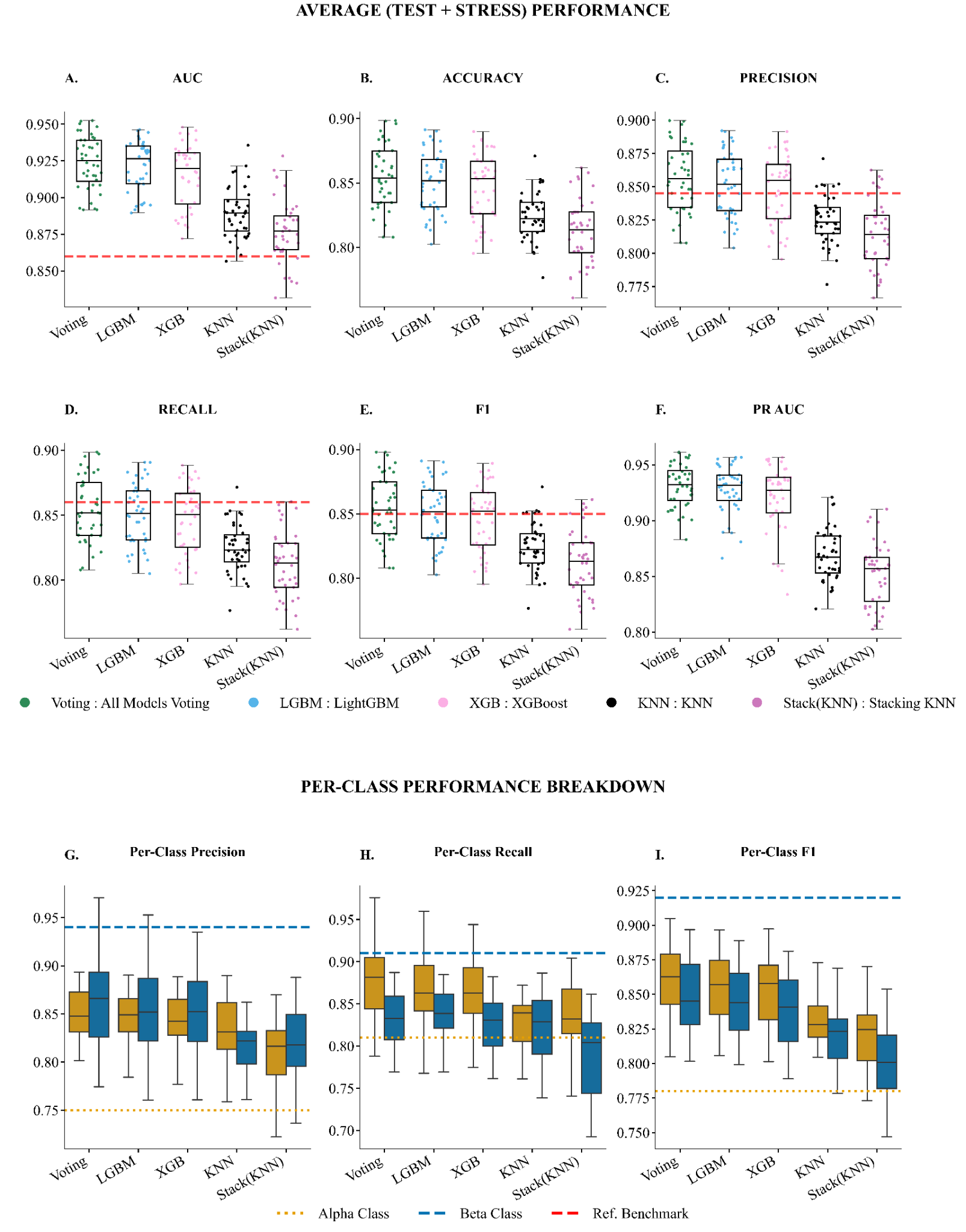
**Supplementary Figure 6 | Aggregated Model Performance Summary Across All Stress Levels.** The figure summarizes the performance of filtered models by aggregating their results across all stress levels and seeds against a pre-defined performance reference across five key evaluation metrics: AUC, Accuracy, F1-Score, Precision, and Recall. The **A-E** panel displays a series of boxplots, each corresponding to a specific metric. Each boxplot illustrates the distribution of performance scores for the models across multiple experimental runs, showing the median, interquartile range, and overall variance. A dashed red line in each plot indicates the reference benchmark for that metric. Similarly, **F-H** panels consist of per-class metric performances: one for Alpha cells (teal) and one for Beta cells (coral). Two distinct horizontal reference lines are shown: a green dotted line for the Alpha cell benchmark ('Ref Alpha') and an orange dashed line for the Beta cell benchmark ('Ref Beta').

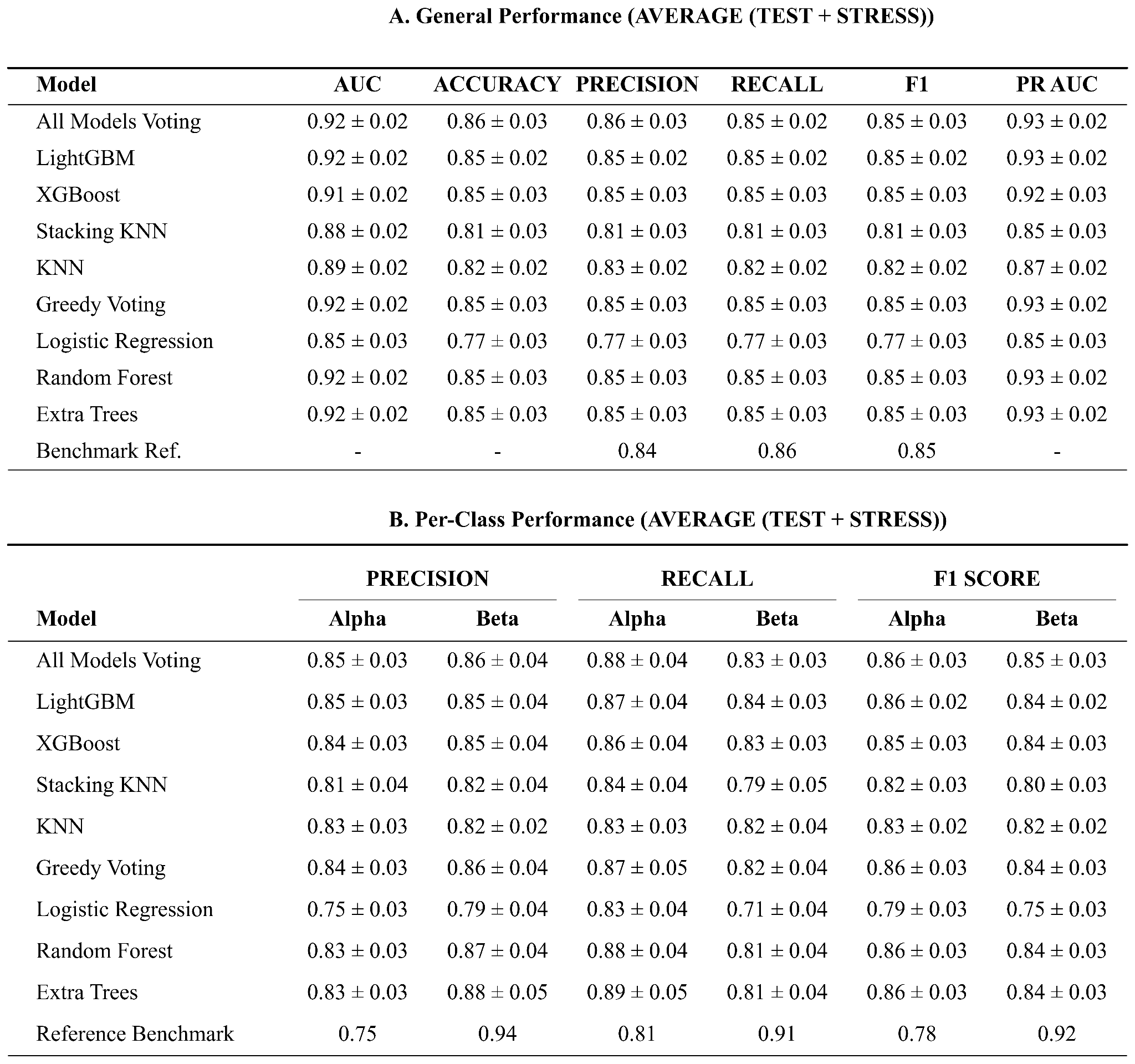
**Supplementary Tables 5 | Aggregated Model Performance Summary Across All Stress Levels.** These tables provide a detailed numerical evaluation of the five distinct machine learning models shown in Supplementary Figure 4. **(A) Global Test Set Results.** This table summarizes the overall performance metrics (AUC, Accuracy, F1-Score, Precision, and Recall). Data is presented as the mean ± standard deviation across experimental runs. The "All models Voting" and "LGBM" classifiers demonstrate top-tier performance, consistently outperforming the Reference values across all key metrics. **(B) Per-Class Metrics.** This table breaks down the performance by specific cell type, detailing Precision, Recall, and F1-Scores for both Alpha and Beta cell populations. This granular view highlights the models' ability to correctly identify and distinguish between the two specific classes relative to the reference standards.

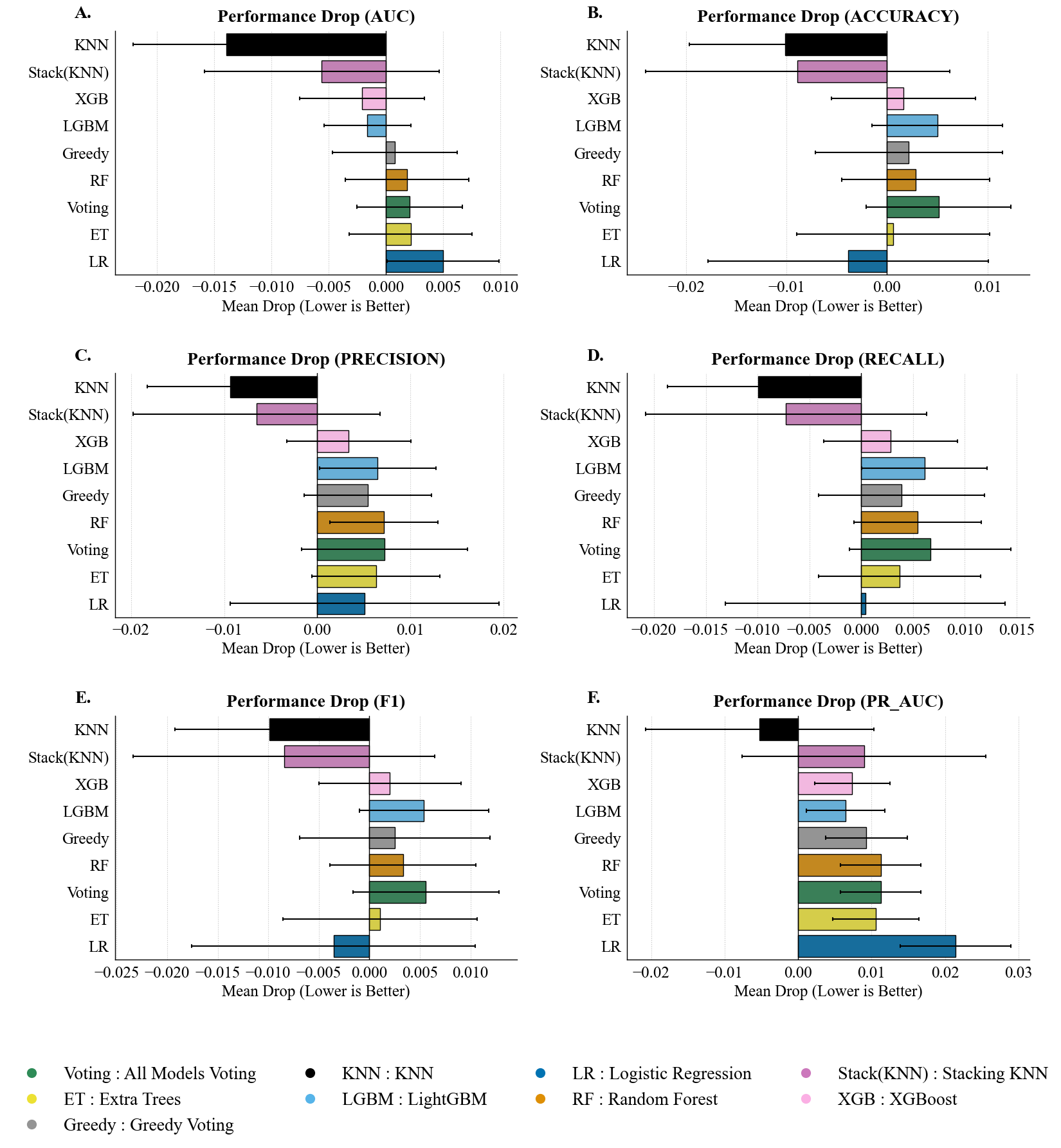
**Supplementary Figure 7 | Drop in Performances of all models across all stress levels for all metrics.** **A-E** The drop is calculated as the difference between a model's performance on the pristine test set (stress level 0) and its average performance across all subsequent stress conditions (stress > 0). The **F** table provides a detailed summary of the mean drop and standard deviation for each model and metric.

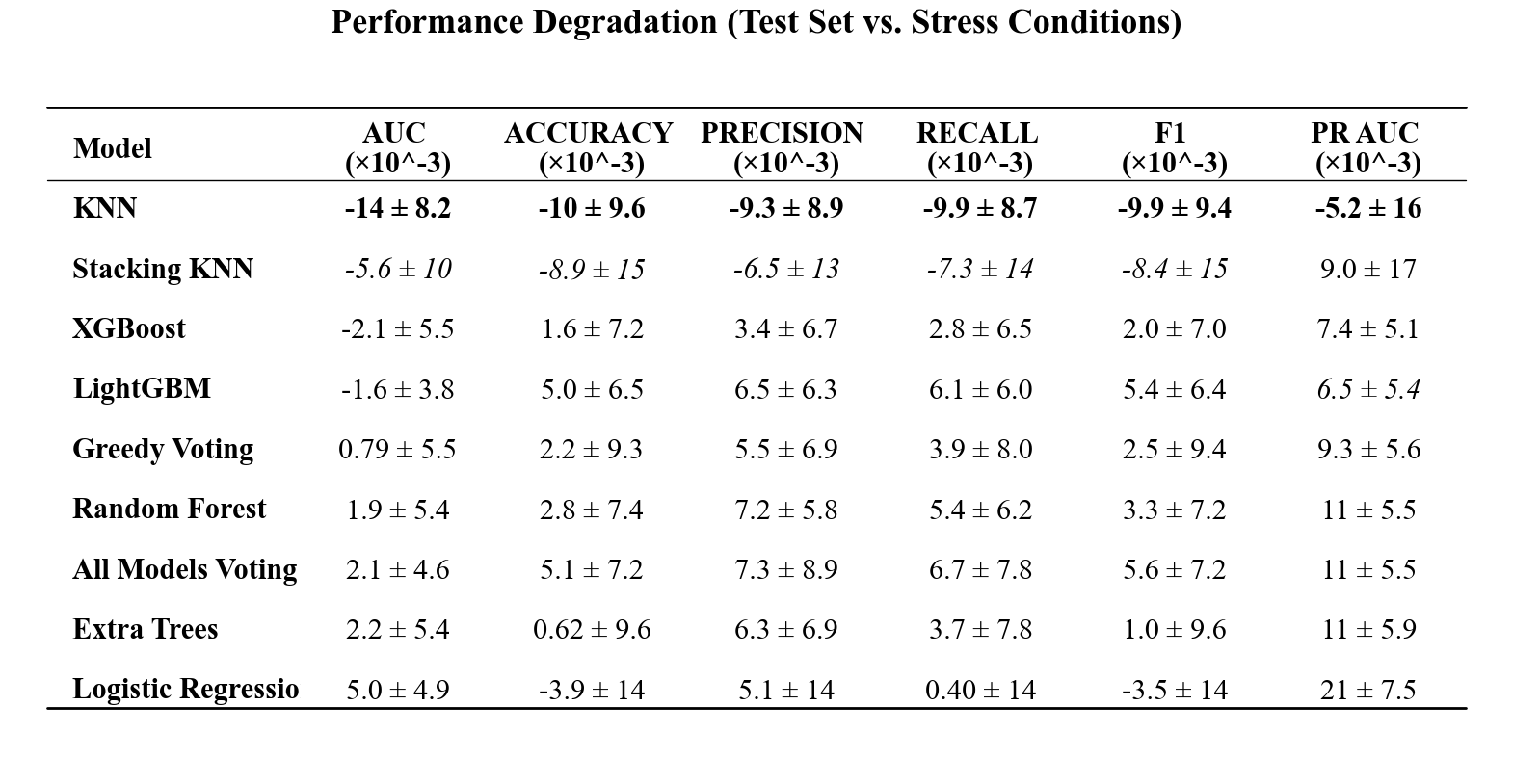

**Supplementary Table 6 | Quantitative Analysis of Model Robustness and Performance Stability.** The table quantifies the impact of geometric stress on model performance by calculating the mean degradation (∆) between the baseline test set and the stress test sets across 10 random seeds. The degradation is defined as ∆ = Test_Score - Stress_Score. Values are reported as Mean 土 Standard Deviation and are scaled by a factor of 10e-3 for readability. To highlight resilience, **bold** values indicate the model with the smallest performance drop (best stability) for a given metric. *Italicized* values indicate the second-best performing model. Negative values imply that the model performance remained stable or slightly exceeded the baseline average under specific stress conditions, indicating high robustness.

Consistent with the findings detailed in the previous section, the LightGBM classifier demonstrates exceptional operational stability across all metrics and stress levels. Performance fluctuations remained negligible; the most pronounced mean reduction was observed in Precision (0.010 ± 0.011), while the Area Under the Curve (AUC) remained virtually static (0.000 ± 0.004). Crucially, no performance decrements reached statistical significance (e.g., AUC drop, *p* = 0.938), suggesting that the model's predictive logic is intrinsically invariant to the applied geometric transformations. This quantitative robustness is visually corroborated by the high density of performance distributions and the remarkably narrow interquartile ranges across conditions (Supplementary Figure 8), confirming the high-dimensional stability of the learned decision boundaries.

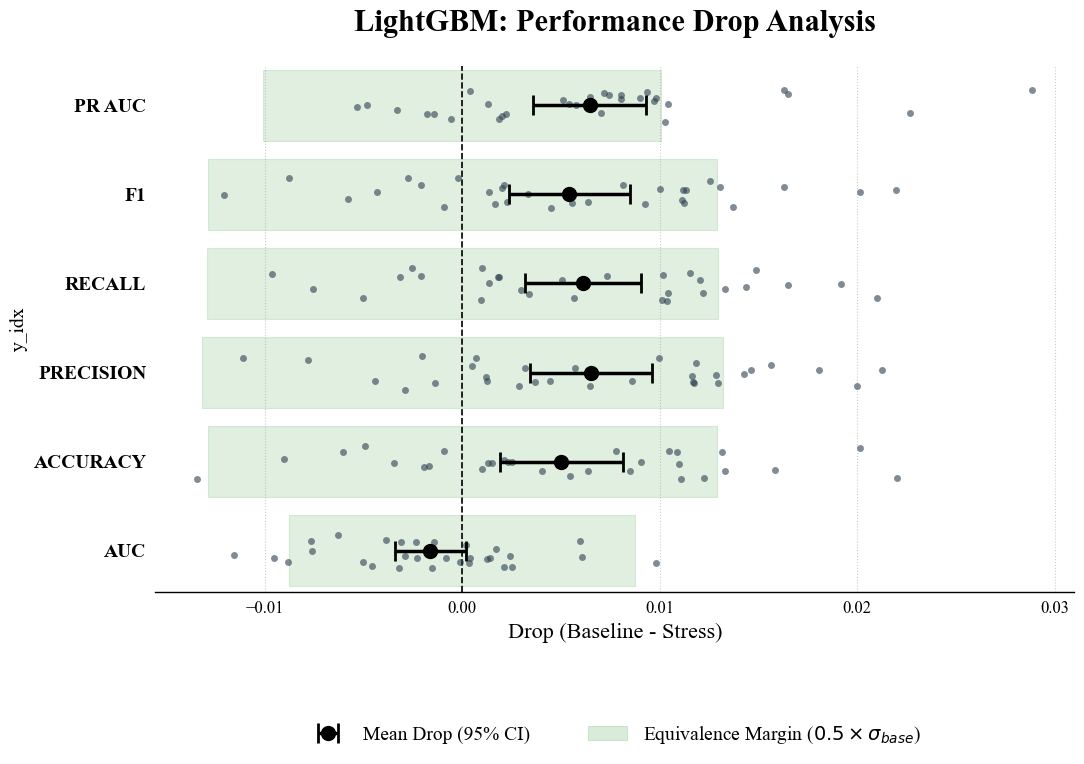
**Supplementary Figure 8 | Statistical Equivalence and Stability Analysis of Model Performance Under Stress.** A Forest plot quantifying the statistical equivalence of the stressed model performances. Black circles represent the mean performance drop (calculated as Baseline minus Stress) averaged across all stress conditions, with error bars denoting the 95% Confidence Interval (CI) of the difference. To rigorously assess structural invariance, a dynamic Equivalence Margin (shaded green band) is established for each metric, defined as half of its baseline standard deviation (0.5 x σ). Individual observations for each seed and stress level are overlaid as dark grey points to illustrate the underlying data distribution (jittered stripplot).

**S2.4** **Feature Importance and Biological Interpretation**

To visualize the decision boundary arising from the complex interactions among discriminative features, we projected the ensemble model’s predicted probabilities onto the first two principal components of the standardized feature space (Supplementary Fig. S9A). The resulting three-dimensional decision surface reveals a pronounced topological asymmetry between the two cell types. α-cells cluster tightly within a low-probability basin, indicating a high degree of morphological homogeneity. In contrast, β-cells span a broad, high-probability plateau, reflecting greater heterogeneity in the organization of intracellular autofluorescence that is nonetheless consistently captured and generalized by the model.

A steep, nearly vertical transition region (“cliff”) separates these two regimes, suggesting that the transition between α- and β-cell autofluorescence pattern phenotypes is abrupt rather than gradual within the learned feature space. This qualitative observation is quantitatively corroborated by the distribution of ensemble prediction probabilities (Supplementary Fig. S9B), which exhibits a strongly bimodal structure. The majority of samples are assigned probabilities close to the extremes (P ≈ 0 or P ≈ 1), while the intermediate region commonly associated with classification ambiguity (0.4 < P < 0.6) is sparsely populated.

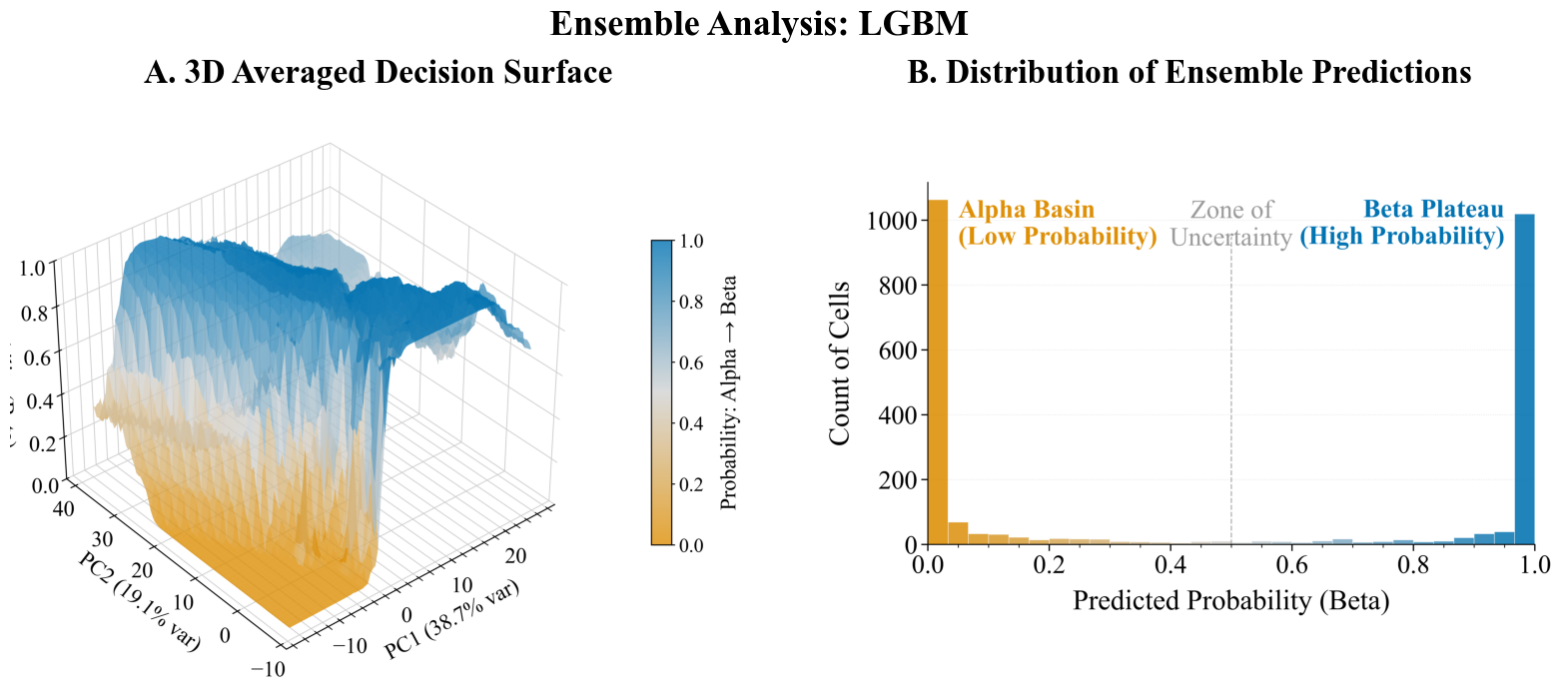

**Supplementary Figure 9 | Topology and confidence of the ensemble decision boundary.** (A) Three-dimensional visualization of the decision surface in the principal component space. The steep vertical transition indicates a sharp separation between class regions. (B) Distribution of predicted probabilities for the test set. The strictly bimodal distribution, with samples clustered at the extremes (0.0 and 1.0) and a negligible "zone of uncertainty," confirms that the ensemble model classifies distinct autofluorescence pattern phenotypes with high confidence and high inter-model agreement.

Together, these results indicate that the ensemble model has learned a robust and well-defined decision boundary, characterized by high inter-model agreement and confident predictions for most samples. The near absence of intermediate probabilities further suggests that morphological ambiguity between α-cells and β-cells is limited in the extracted feature space, reinforcing the biological relevance and discriminative power of the selected descriptors. Notably, given the spectral window employed in this study, the structured autofluorescence patterns captured by the model are consistent with differences in lipofuscin granule abundance and spatial distribution between α- and β-cells. The sharp decision boundary observed in the reduced feature space, therefore, likely reflects biologically grounded differences in intracellular autofluorescent organization, supporting the interpretation that cell identity is structurally encoded in these intensity patterns rather than arising from abstract feature separability.

**S2.5**	**On the importance of Augmentation**

Although the extracted feature set was explicitly engineered to be invariant to expected real-world transformations, we observed that synthetic data augmentation can nonetheless introduce small, non-ideal perturbations in the feature space. This effect arises primarily from interpolation inherent in geometric transformations, particularly image rotations at angles that are not divisible by 90°. Such fractional pixel shifts produce subtle variations in pixel intensities, which in turn propagate to the computed feature vectors, even for descriptors designed to be invariant. These augmentation-induced deviations are illustrated in Supplementary Fig. 10.

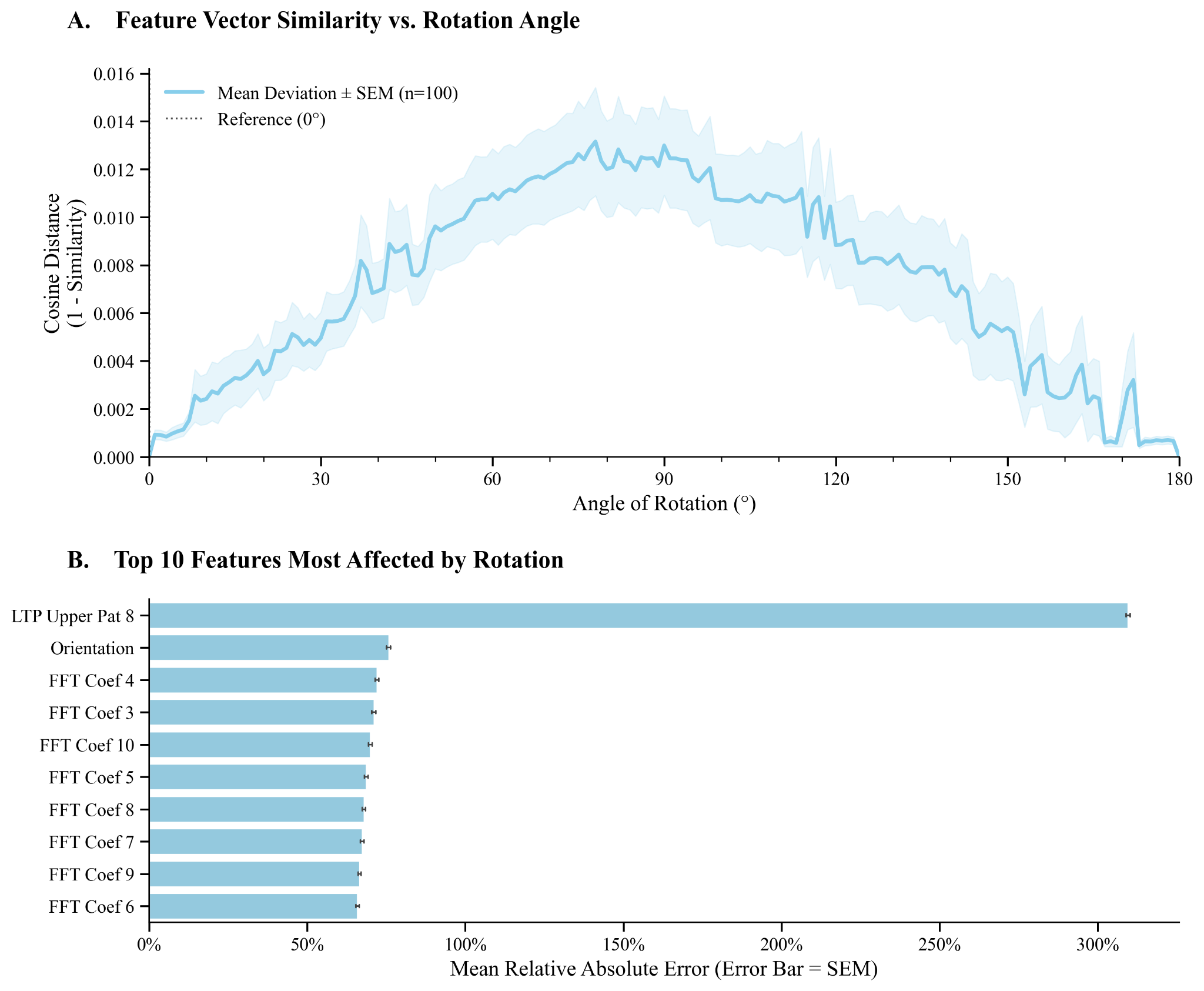

**Supplementary Figure 10 | Statistical Proof of Interpolation-Induced Feature Deviation.** Panel A) Feature Vector Similarity vs. Rotation Angle. This panel illustrates the relationship between cosine distance and the angle of rotation for feature vectors, based on 100 images and 95% confidence intervals. Panel B) Top 10 Features Most Affected by Rotation (Relative Error). This panel identifies the ten features that exhibit the highest mean relative absolute error under rotational augmentation, sorted in descending order of error.

To quantitatively assess the contribution of data augmentation to model robustness, we conducted a controlled experiment in which the entire training and evaluation pipeline was repeated under identical conditions, with all augmentation steps removed—except for those applied during the designated stress-test phase. Under this configuration, a pronounced degradation in model performance was observed during stress testing. Notably, when the standard filtering procedure was retained, none of the trained models met the predefined performance criteria required to pass the stress test. As a result, the filtering step was removed in subsequent analyses to enable a complete and unbiased comparison across models. Performance metrics for all evaluated models under stress-test conditions are reported in Supplementary Fig. S11 and Supplementary Table 7 (for brevity, only stress level 3 is shown).

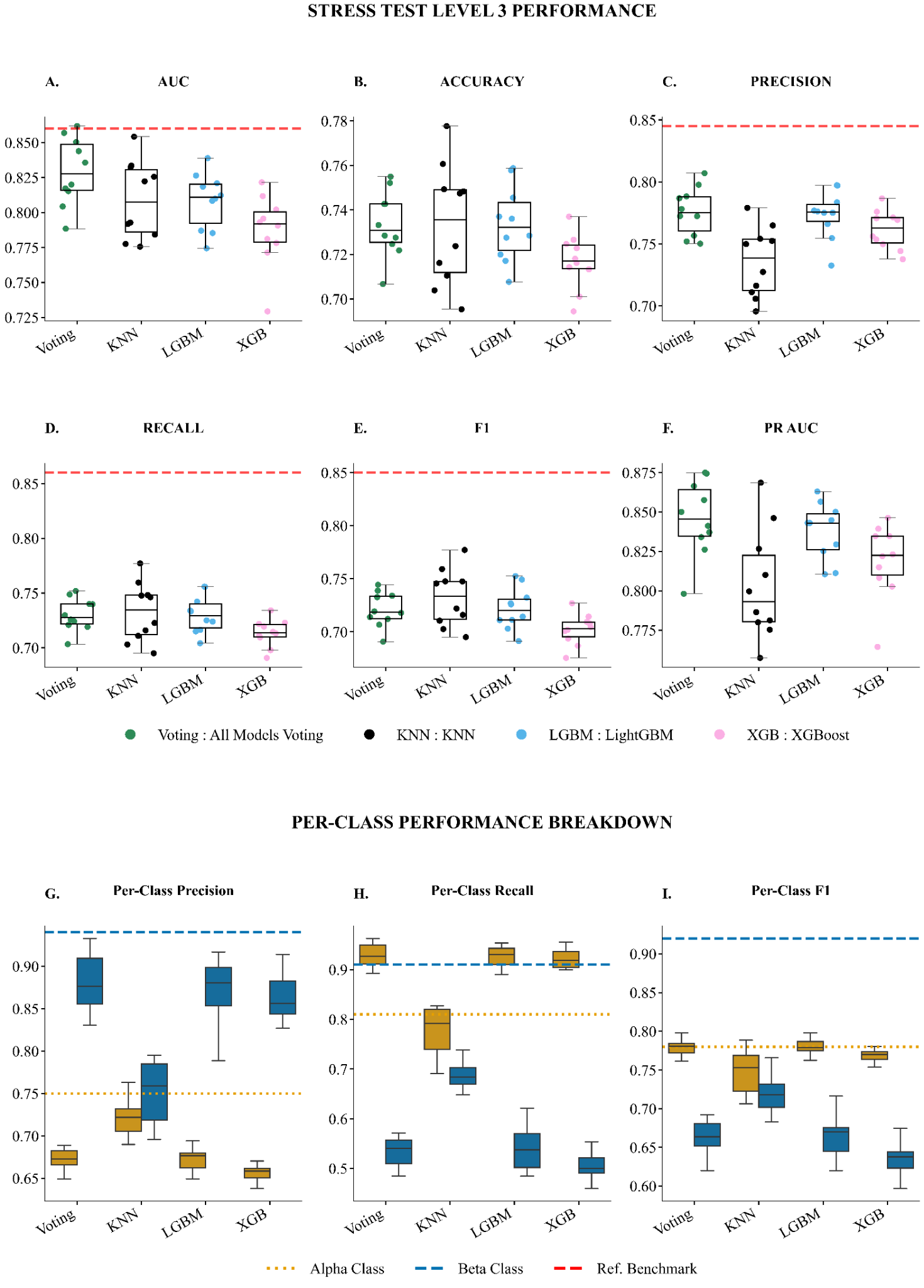
**Supplementary Figure 11| Comparative Performance of Machine Learning Models on the Stress Test Level 3.** The figure presents a comprehensive comparison of five distinct machine learning models against a pre-defined performance reference across five key evaluation metrics: AUC, Accuracy, F1-Score, Precision, and Recall. **A-E** panel displays a series of boxplots, each corresponding to a specific metric. Each boxplot illustrates the distribution of performance scores for the models across multiple experimental runs, showing the median, interquartile range, and overall variance. A dashed red line in each plot indicates the reference benchmark for that metric. Similarly, **F-H** panels consist of per-class metric performances: one for Alpha cells (teal) and one for Beta cells (coral). Two distinct horizontal reference lines are shown: a green dotted line for the Alpha cell benchmark ('Ref Alpha') and an orange dashed line for the Beta cell benchmark ('Ref Beta').

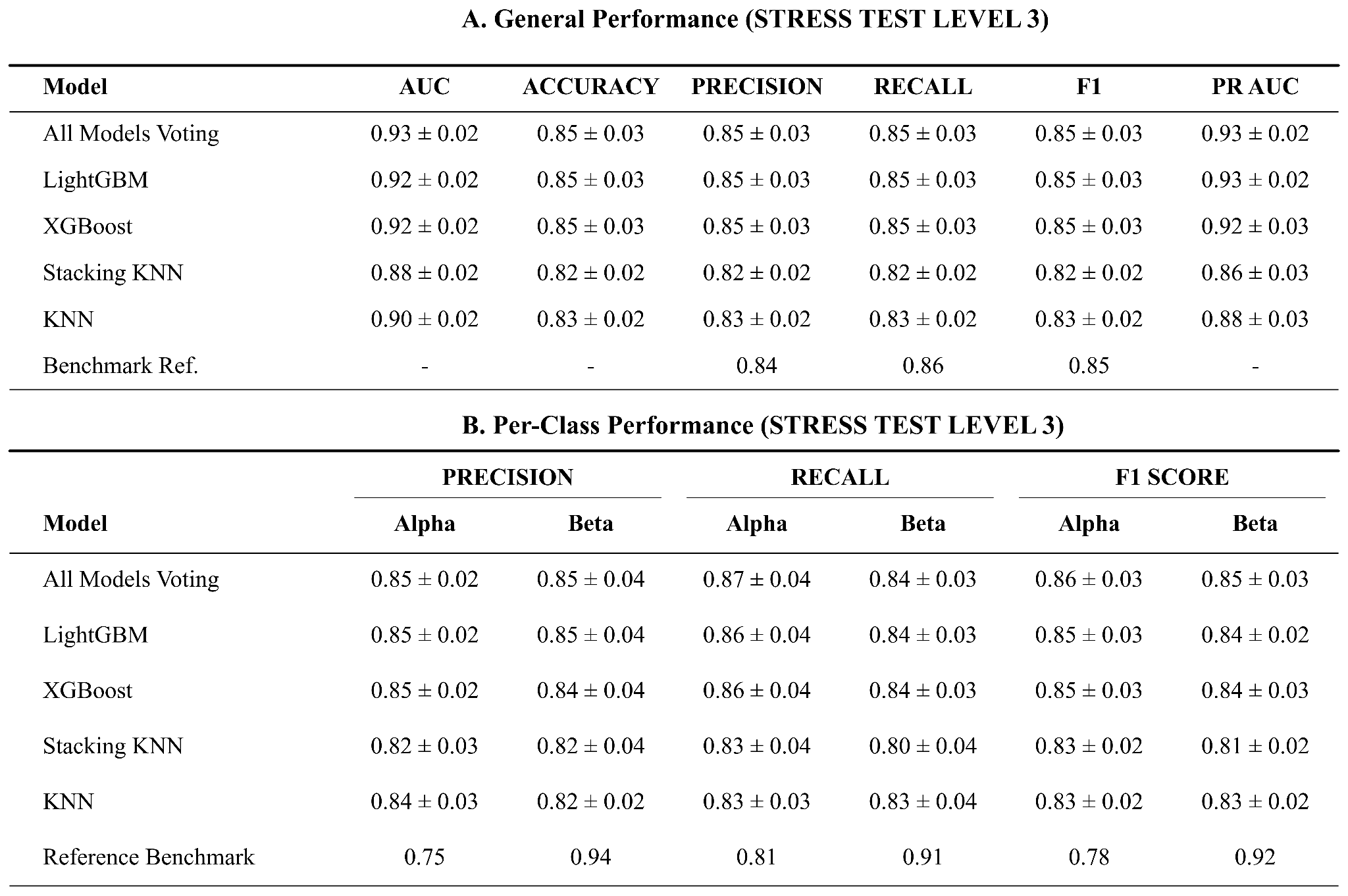

**Supplementary Table 7 | Model Performance Summary on Stress Test Levels 3.** These tables provide a detailed numerical evaluation of the five distinct machine learning models shown in Supplementary Figure 8. (A) Global Test Set Results. This table summarizes the overall performance metrics (AUC, Accuracy, F1-Score, Precision, and Recall). Data is presented as the mean ± standard deviation across experimental runs. (B) Per-Class Metrics. This table breaks down the performance by specific cell type, detailing Precision, Recall, and F1-Scores for both Alpha and Beta cell populations.

Across all models, both AUC and accuracy exhibit substantial reductions relative to test-set performance. For example, the Greedy Voting ensemble shows a decrease in AUC from 0.93 on the test set to 0.86 and 0.83 under increasing stress levels. However, the most pronounced degradation is observed in the F1 score and recall metrics. Compared to the test set, stress-test evaluations show drops of 0.10 or greater, accompanied by severe class-specific imbalance. In the case of Greedy Voting, recall for α-cells and β-cells shifts from 0.90 and 0.82 on the test set to 0.91 and 0.59, respectively, at the third stress level, indicating a marked loss of sensitivity for the β-cell class.

This trend is summarized in Supplementary Fig. S12 (and detailed in Supplementary Table 8), which reports the average performance degradation at the highest stress level across all evaluated models. All classifiers exhibit measurable declines, with linear models showing the most pronounced reductions across metrics, particularly in AUC and F1 score. Ensemble-based approaches display comparatively smaller performance losses, indicating increased robustness to stress-induced distributional shifts.

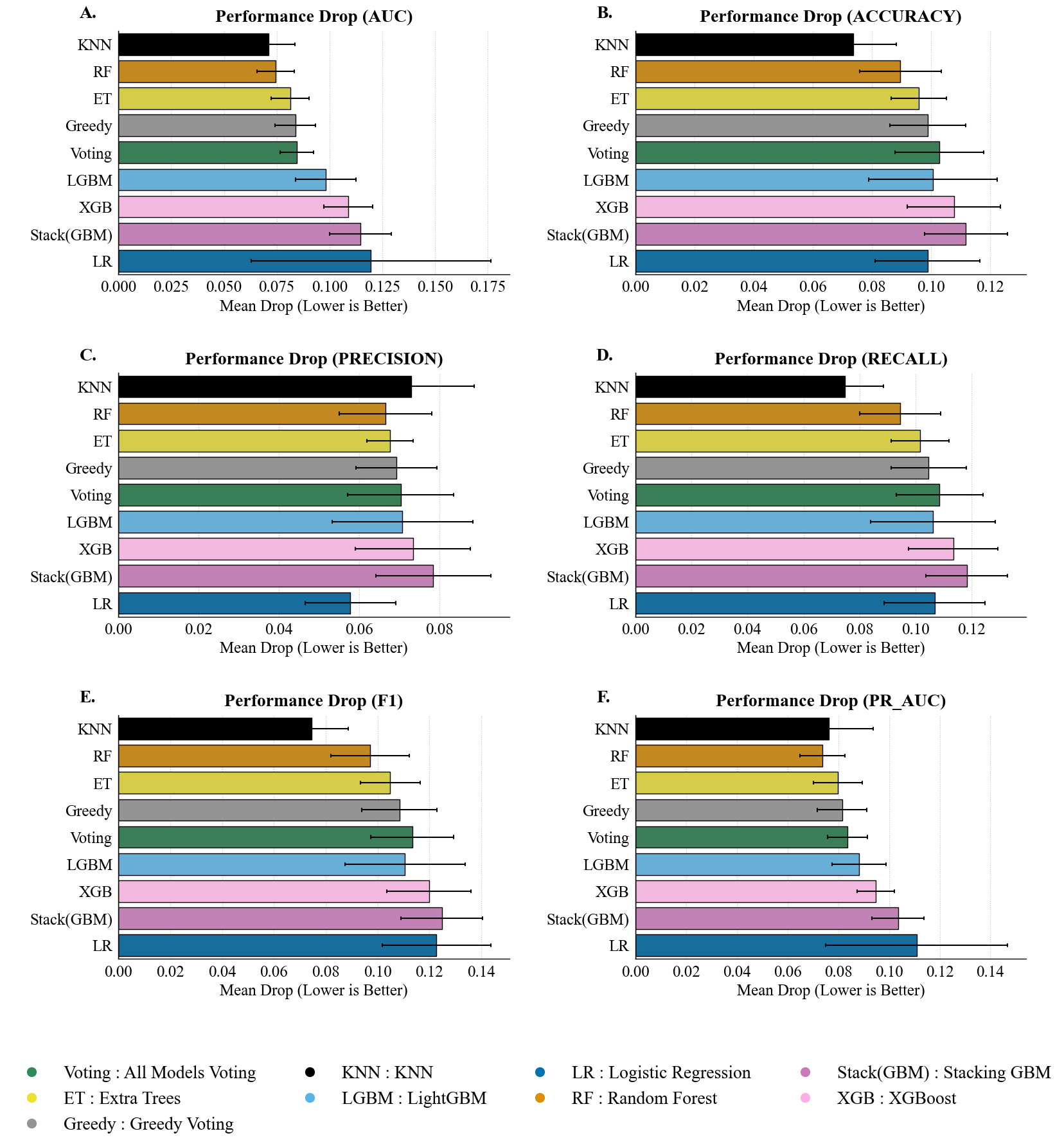

**Supplementary Figure 12 | Drop in Performances of all models across all stress levels for all metrics.** **A-E** The drop is calculated as the difference between a model's performance on the pristine test set (stress level 0) and its average performance across all subsequent stress conditions (stress > 0). The **F** table provides a detailed summary of the mean drop and standard deviation for each model and metric.

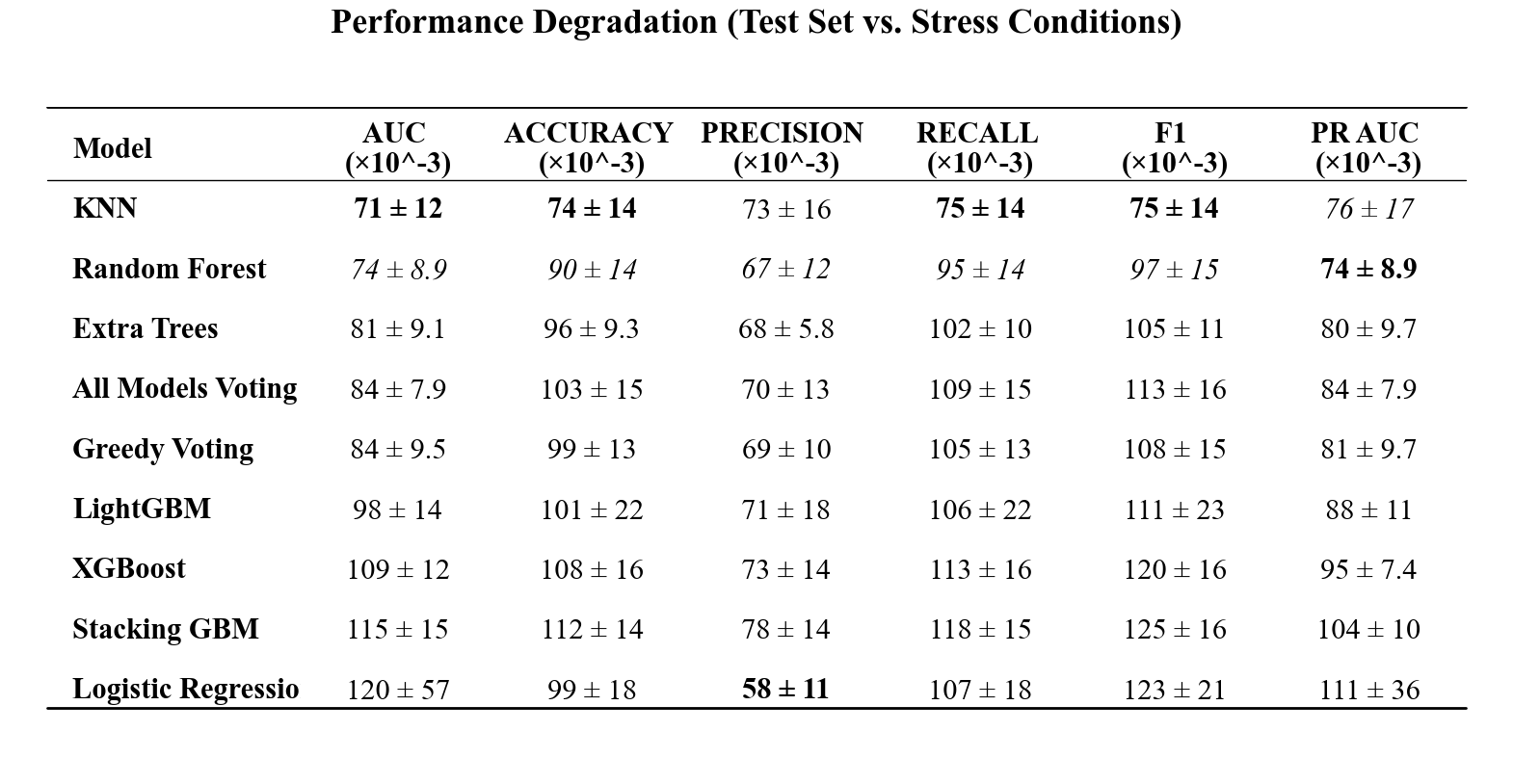
**Supplementary Table 8 | Quantitative Analysis of Model Robustness and Performance Stability.** The table quantifies the impact of geometric stress on model performance by calculating the mean degradation (∆) between the baseline test set and the stress test sets across 10 random seeds. The degradation is defined as ∆ = Test_Score - Stress_Score. Values are reported as Mean 土 Standard Deviation and are scaled by a factor of 10e-3 for readability. To highlight resilience, **bold** values indicate the model with the smallest performance drop (best stability) for a given metric. *Italicized* values indicate the second-best performing model. Negative values imply that the model performance remained stable or slightly exceeded the baseline average under specific stress conditions, indicating high robustness.

Nevertheless, even the most resilient models experience non-negligible decreases in recall and F1 score, reflecting a systematic loss of class-specific sensitivity under stress. The observed variability in degradation across models further highlights the instability of classifier behavior when exposed to distributional perturbations. Taken together, these results demonstrate that while ensembling partially mitigates robustness loss, no evaluated model remains invariant to stress-test conditions, underscoring the importance of robustness assessment beyond standard test-set evaluation.

**S2.6**	**Justification for the Inclusion of Extended α Cell Data**

Beyond improving robustness to real-world variability, a central objective of this study was to enhance the model’s generalization capacity by enabling it to process a broader and more diverse set of images than previously possible. To this end, a substantial fraction of the original dataset—primarily α-cell images—had been excluded in earlier processing stages. It was therefore critical to assess whether reintroducing a subset of these previously eliminated samples (Dataset++) contributed meaningful discriminative information rather than introducing noise.

To evaluate this, we compared models trained on the extended dataset (Dataset++) with those trained on the restricted dataset (Dataset) under two class-balancing strategies: standard data augmentation and SMOTE. Statistical significance was assessed using the Wilcoxon signed-rank test (α = 0.01) across 10 independent random seeds. Model performance was evaluated using two complementary analytical frameworks: Intra-Dataset Performance and Inter-Dataset Performance.

Intra-dataset analysis measured performance stability across different train–test splits derived from the same dataset, thereby isolating sensitivity to internal data variability. In contrast, inter-dataset analysis directly assessed generalization by evaluating models trained on a source dataset against a distinct, independent target dataset, providing a stringent test of robustness to distributional shifts. Results are summarized in Supplementary Table S4 and Supplementary Fig. S13.

The inter-dataset performance analysis demonstrates a clear advantage for models trained on Dataset++ relative to those trained on the restricted Dataset, even when the latter is balanced using standard augmentation techniques (Supplementary Table 9). This indicates that the additional images in Dataset++ contribute genuinely informative and discriminative content that cannot be reproduced through geometric transformations alone.

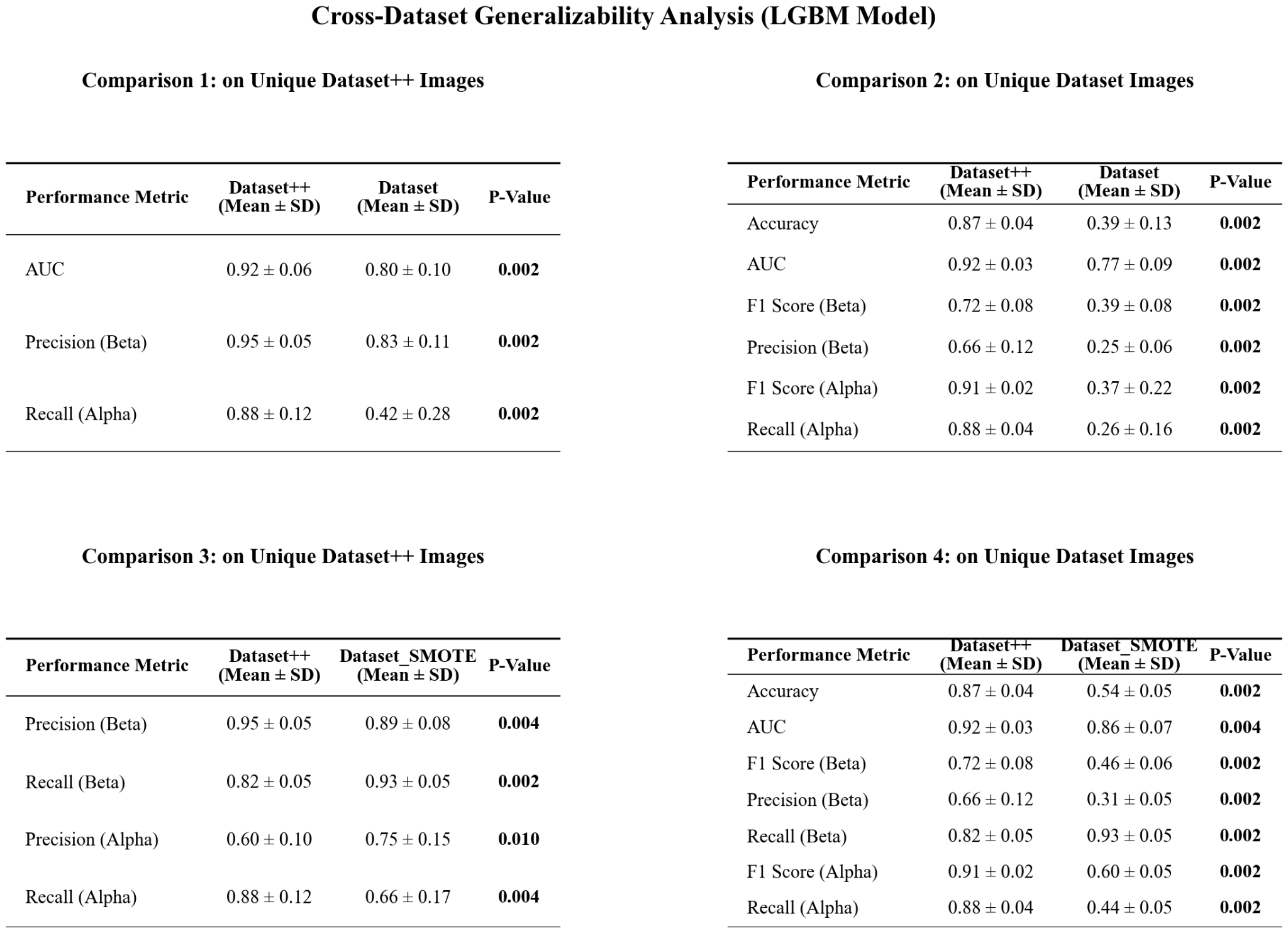

**Supplementary Table 9 |** **Cross-Dataset Generalizability Analysis of LGBM Models.** This figure presents a comprehensive evaluation of the generalizability of the developed LightGBM (LGBM) models by assessing their performance on unseen test data from different source datasets. The four tables display performance metrics (Mean ± Standard Deviation) for pairwise comparisons between the Dataset, Dataset++, and Dataset_SMOTE models. The top row presents results on the Unique Dataset++ Test images, comparing the Dataset++ model against the Dataset model (left) and the Dataset_SMOTE model (right). The bottom row presents results on the Unique Dataset Test images, comparing the Dataset model (left) and the Dataset_SMOTE model (right) against the Dataset++ model as a reference. Statistical significance was assessed using the Wilcoxon signed-rank test, with ** indicating a p-value < 0.01. For each performance metric, the significantly higher mean value between the two compared models is highlighted in bold.

To disentangle the effect of increased informational content from that of class rebalancing, we conducted a direct comparison between models trained on Dataset++ and models trained on the restricted Dataset balanced using SMOTE. Across all major performance metrics, the Dataset++ models significantly outperformed their SMOTE-based counterparts. Specifically, Dataset++ achieved higher Accuracy (0.87 ± 0.04 vs. 0.54 ± 0.05), AUC (0.92 ± 0.03 vs. 0.86 ± 0.07), and β-cell F1 score (0.72 ± 0.08 vs. 0.46 ± 0.06). A class-specific analysis reveals trade-offs characteristic of synthetic oversampling. While the Dataset++ model exhibits superior generalization, reflected in significantly higher α-cell recall and β-cell precision, the SMOTE-balanced model achieves a higher β-cell recall (0.93 ± 0.05 vs. 0.82 ± 0.05) at the expense of β-cell precision (0.31 ± 0.05 vs. 0.66 ± 0.11). This pattern indicates that SMOTE expands minority-class decision boundaries aggressively, capturing more positive samples but introducing noise that substantially increases false positives. The most critical insight emerges from the analysis of per-class precision under Stress Level 3 conditions (Supplementary Fig. 11). Models trained on Dataset++ maintain balanced and stable precision across both the test set and stress-test conditions. In contrast, the SMOTE-trained model exhibits pronounced instability in its class-specific behavior. On the standard test set, the SMOTE model shows low α-cell precision (0.73 ± 0.05) and high β-cell precision (0.90 ± 0.02); however, under stress-test conditions, this pattern reverses sharply, with α-cell precision increasing to 0.90 ± 0.02 and β-cell precision dropping to 0.75 ± 0.05.

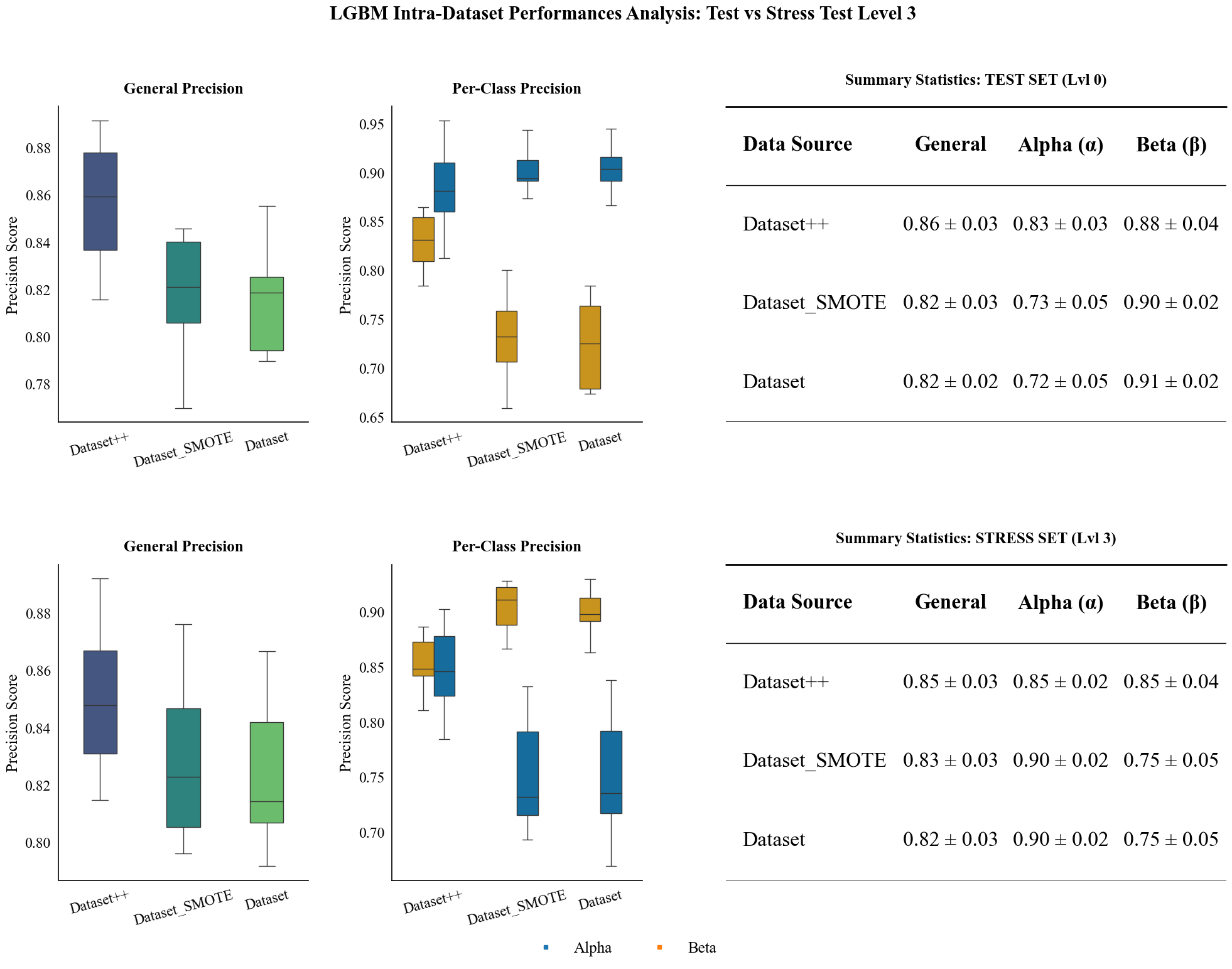

**Supplementary Figure 13 | Intra-Dataset Precision Analysis: Comparison between Standard Test and Stress Level 3**. This figure illustrates the robustness of the LightGBM models trained on different data configurations (Dataset, Dataset++, and Dataset_SMOTE) when subjected to high-intensity stress conditions (Level 3 noise injection). The top row displays the overall precision distributions for the standard test set (Left) versus Stress Level 3 (Center), accompanied by a summary table (Right) detailing the Mean ± Standard Deviation for each data source. The bottom row breaks down these performances by class (Alpha vs. Beta), showing the class-specific precision distributions for the standard test set (Left) and Stress Level 3 (Center), with a corresponding summary table (Right) provided. The boxplots visualize the median, interquartile range, and outliers across 10 random seeds, demonstrating the impact of stress on model stability and class-wise performance.

**S2.7 Pipeline Speed and Computational Efficiency Analysis**

An initial component-level profiling analysis (Supplementary Figure 14) indicated that feature extraction constitutes the primary computational cost, accounting for more than 90% of total processing time. Under optimized batch-processing conditions, in which large numbers of images are evaluated concurrently, the system exhibited efficient performance. The serial implementation achieved a mean per-image processing time of 0.74 ± 0.05 ms, while the parallel implementation reduced this to 0.35 ± 0.04 ms, corresponding to an effective throughput of approximately 2,800 images per second.

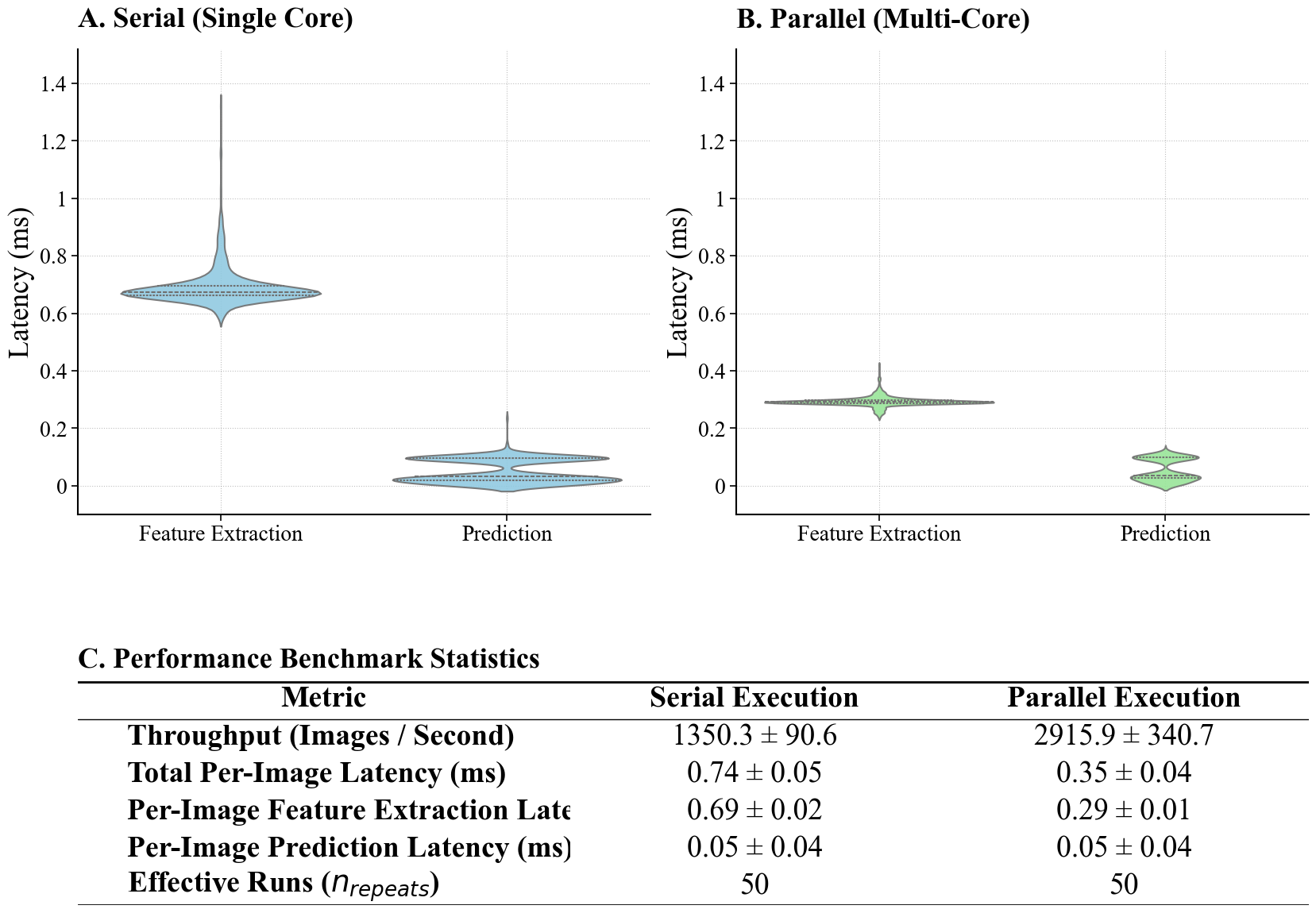

**Supplementary Figure 14 | Aggregated Pipeline Performance Dashboard.** This figure presents a comparative performance analysis of the feature extraction and prediction pipeline under serial (single-core) and parallel (multi-core) execution modes. (Top) Violin plots illustrate the distribution of latency (ms) for both feature extraction and prediction stages. The blue violins represent serial execution, showing higher variability and mean latency, particularly for feature extraction, while the green violins depict the improved efficiency and tighter distribution of parallel execution. (Bottom) A summary table quantifies the performance metrics, detailing the mean throughput (Images/Second) and per-image latency (ms) with standard deviations for both execution modes, based on 50 effective runs.

Stress testing across a range of input sizes (Supplementary Figure 15) showed that serial and parallel implementations perform similarly under low-load conditions (N < 100). At higher loads, however, the parallel implementation demonstrated more favorable scaling behavior, maintaining stable throughput as input size increased, whereas the serial implementation exhibited saturation effects.

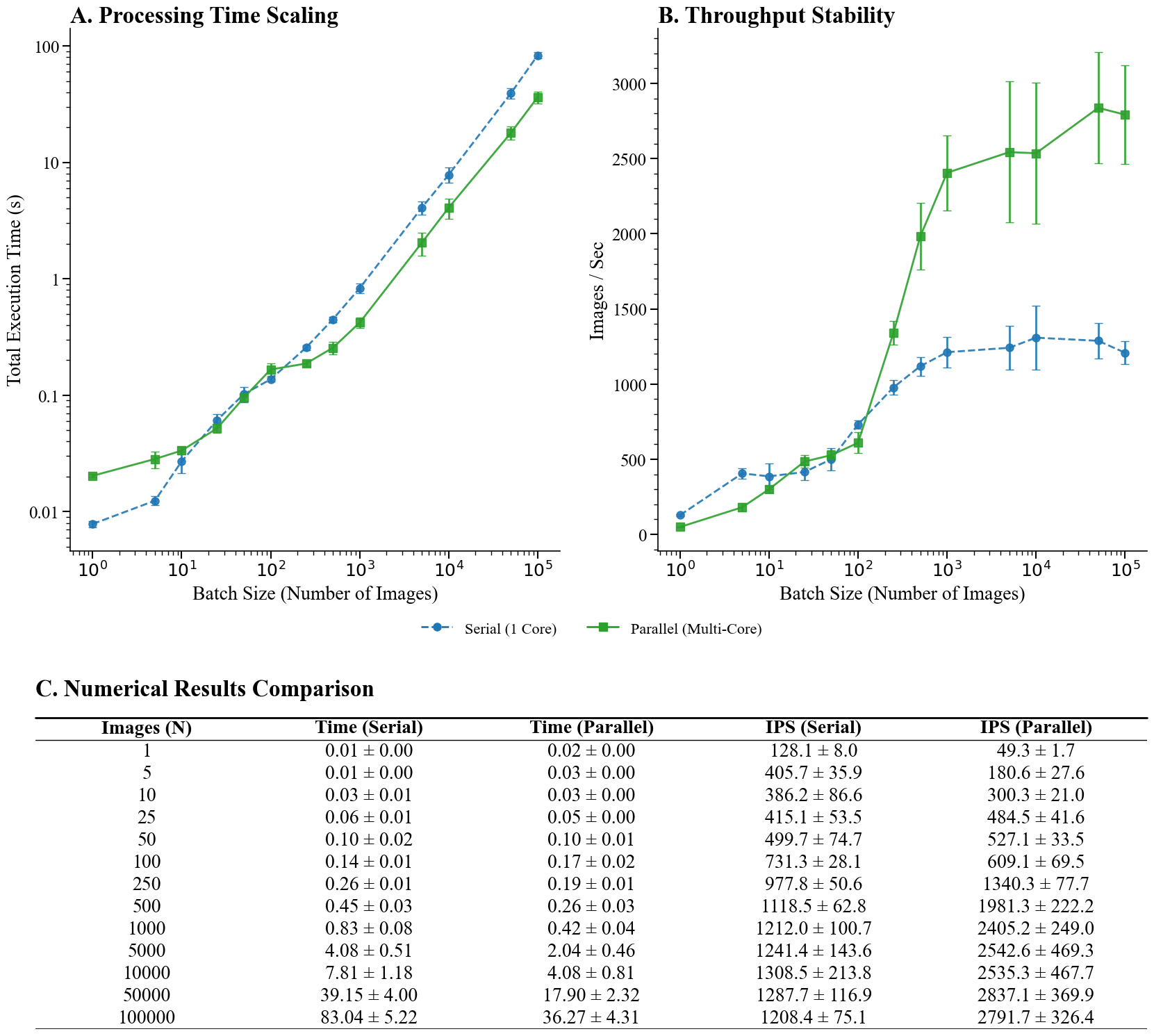

**Supplementary Figure 15 | Scalability Stress Test and Throughput Stability.** This figure evaluates the system's performance scaling across varying batch loads ranging from 1 to 100’000 images. **A.** The log-log plot demonstrates the total execution time scaling, revealing a crossover point at approximately *N=100* where parallel execution (green solid line) begins to outperform serial execution (blue dashed line) as initialization overheads are amortized. **B.** The throughput stability analysis highlights the saturation limits of both architectures, with the parallel engine reaching a higher steady-state capacity of approximately 2,800 Images/Sec compared to the serial engine's limit of roughly 1,200 Images/Sec. **C.** The tabulated numerical results provide the precise mean values and standard deviations for execution time and throughput at each tested batch interval.

A different performance profile emerged when evaluating single-frame processing, which more closely reflects the operational requirements of a live camera stream. As illustrated in Supplementary Figure 16, the legacy pipeline exhibited a median latency of 17 ms per frame, substantially higher than the per-image latency observed under batch-processing conditions. This discrepancy arises from the inability to amortize fixed overheads—such as data structure initialization and function call setup—when processing individual frames. In batch mode, these overheads are distributed across many samples and therefore have a limited impact, whereas in a single-input, single-output execution model, they are incurred for every frame.

To address this limitation, a specialized inference pipeline was developed with a streamlined data flow and reduced reliance on heavyweight abstractions. This redesign significantly reduced per-frame overhead, lowering the median end-to-end latency to 4.2 ms, representing and achieving a frame rate of 209 FPS. Importantly, these performance improvements were achieved without compromising numerical correctness. The optimized pipeline produced identical outputs to the original implementation, yielding a mean absolute error of 0.00 and complete label agreement. Together, these results indicate that the revised architecture effectively reconciles computational efficiency with real-time operational constraints, enabling practical deployment in latency-sensitive applications.

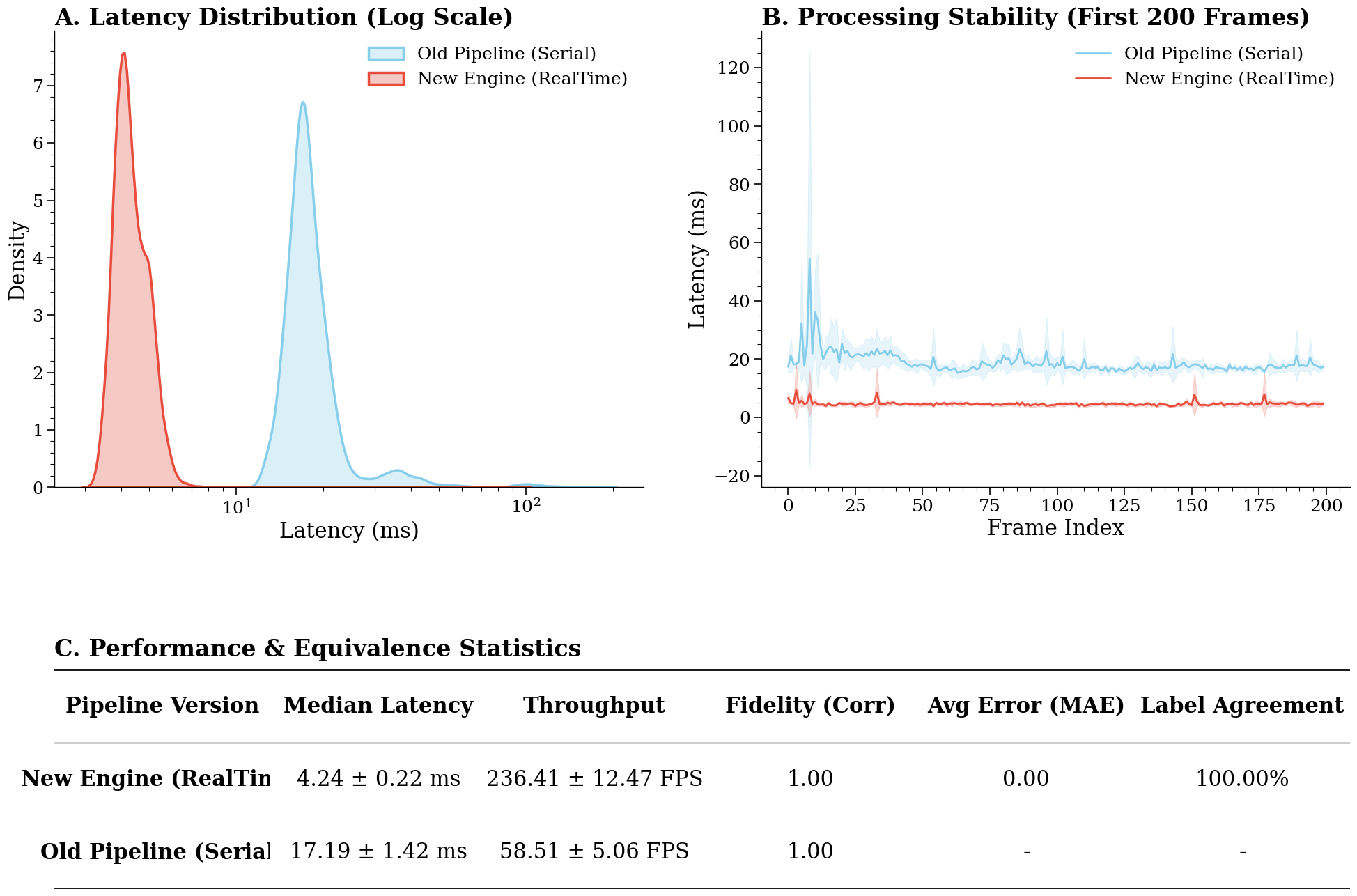
**Supplementary Figure 16 | Real-Time Feasibility and Comparative Validation.** This dashboard contrasts the performance of the legacy serial pipeline (red) against the optimized real-time engine (green) under single-frame streaming conditions. A. The Kernel Density Estimation (KDE) plot on a logarithmic scale reveals a significant shift in latency distribution, with the new engine centering around 4.8 ms compared to the legacy pipeline’s 19 ms, indicating a substantial reduction in processing delay. B. The temporal stability analysis over the first 200 frames demonstrates the reduction in processing jitter and the elimination of per-frame initialization overheads in the new engine. C. The statistical summary confirms a nearly four-fold increase in throughput (~210 FPS vs. ~53 FPS) while maintaining 100% label agreement and zero numerical error (MAE = 0.00), verifying that performance gains were achieved without compromising predictive fidelity.

Altogether, these results indicate that the pipeline is sufficiently flexible to elaborate large datasets of images and maintain a high throughput of sequential images, in our test scenario. This can optimistically indicate that future real-time setup and cloud-based classification may be feasible, despite the challenges that the true operational environment will naturally present.
